# Modulation of dopamine D_1_ receptors via histamine H_3_ receptors is a novel therapeutic target for Huntington’s disease

**DOI:** 10.1101/837658

**Authors:** David Moreno-Delgado, Mar Puigdellívol, Estefanía Moreno, Mar Rodríguez-Ruiz, Joaquín Botta, Paola Gasperini, Anna Chiarlone, Lesley A. Howell, Marco Scarselli, Vicent Casadó, Antoni Cortés, Sergi Ferré, Manuel Guzmán, Carme Lluís, Jordi Alberch, Enric Canela, Sílvia Ginés, Peter J. McCormick

## Abstract

Early Huntington’s disease (HD) include over-activation of dopamine D_1_ receptors (D_1_R), producing an imbalance in dopaminergic neurotransmission and cell death. To reduce D_1_R over-activation, we present a strategy based on targeting complexes of D_1_R and histamine H_3_ receptors (H_3_R). Using an HD striatal cell model and HD organotypic brain slices we found that D_1_R-induced cell death signaling and neuronal degeneration, are mitigated by an H_3_R antagonist. We demonstrate that the D_1_R-H_3_R heteromer is expressed in HD animal models at early but not late stages of HD, correlating with HD progression. In accordance, we found this target expressed in human control subjects and low-grade HD patients. Finally, treatment of HD mice with an H_3_R antagonist prevented cognitive and motor learning deficits, as well as the loss of heteromer expression. Taken together, our results indicate that D_1_R - H_3_R heteromers play a pivotal role in dopamine signaling and represent novel targets for treating HD.

**Impact Statement:** Progression of Huntington’s disease can be slowed by altering dopamine signalling through the Dopamine 1 receptor - Histamine 3 receptor heteromer.

## Introduction

Huntington’s disease (HD) is a dominant inherited progressive neurodegenerative disorder caused by expansion of a CAG repeat, coding a polyglutamine repeat within the *N*-terminal region of huntingtin protein (HDCRG, 1993; Vonsattel and DiFiglia, 1998). Although dysfunction and death of striatal medium-sized spiny neurons (MSSNs) is a key neuropathological hallmark of HD(Ferrante et al., 1991; Vonsattel et al., 1985), cognitive deficits appear long before the onset of motor disturbances (Lawrence et al., 2000; Lemiere et al., 2004). It has been postulated that alterations in the dopaminergic system may contribute to HD neuropathology (Chen et al., 2013; Jakel and Maragos, 2000), as dopamine (DA) plays a key role in the control of coordinated movements. Increased DA levels and DA signaling occur at early stages of the disease(Chen et al., 2013; Garret et al., 1992; Jakel and Maragos, 2000), resulting in an imbalance in striatal neurotransmission initiating signaling cascades that may contribute to striatal cell death(Paoletti et al., 2008; Ross and Tabrizi, 2011). Several studies demonstrated that DA receptor antagonists and agents that decrease DA content reduce chorea and motor symptoms while dopaminergic stimulation exacerbate such symptoms(Huntington Study Group, 2006; Mestre et al., 2009; Tang et al., 2007).

Within the striatum, two different MSSNs populations can be distinguished: 1) MSSNs expressing enkephalin and dopamine D_2_ receptors (D_2_R), which give rise to the indirect striatal efferent pathway, and 2) MSSNs expressing substance P and dopamine D_1_ receptors (D_1_R), comprising the direct striatal efferent pathway. Recently, several studies with experimental models have changed the traditional view that D_2_R-MSSNs are more vulnerable in HD(Cepeda et al., 2008; Kreitzer and Malenka, 2007), proposing a new view in which D_1_R-MSSNs are more vulnerable to the HD mutation. In this view, it has been demonstrated that mutant huntingtin enhances striatal cell death through the activation of D_1_R but not D_2_R (Paoletti et al., 2008). More recently, it has been described that, at early stages of the disease, HD mice show an increase in glutamate release onto D_1_R neurons but not D_2_R neurons while, later in the disease, glutamate release is selectively decreased to D_1_R cells (Andre et al., 2011), indicating that several changes occur in D_1_R neurons at both early and late disease stages. Strategies that might reduce D_1_R signaling could prove successful towards preventing HD (10;14;17;18). However, D_1_Rs are highly expressed in many tissues (19) and broad use of D_1_R antagonists as a preventive treatment has important drawbacks including locomotor impairments (20), or induce depression, parkinsonism and sedation in HD patients (12;21).

Histamine is an important neuromodulator with four known G protein-coupled receptors (GPCRs). H_3_Rs are expressed in brain regions involved in both motor function (striatum) and cognition, such as the cortex, thalamus, hypothalamus, hippocampus and amygdala (22). It is known that in at least striatal GABAergic dynorphinergic neurons (23–25), both D_1_R and H_3_R are co-expressed and we and others have found that they establish functional negative interactions by forming molecular complexes termed heteromers (26;27). Hence, in this work, we hypothesized that targeting D_1_R through these receptor complexes of D_1_R and H_3_R might serve as a more efficient and targeted strategy to slow the progression of HD. Specifically, we demonstrate that D_1_R-H_3_R heteromers are expressed and functional in early HD stages but are lost in late stages. An H_3_R antagonist acting through D_1_R-H_3_R heteromers acts as a protective agent against dopaminergic imbalance in early HD stages improving learning and long-term memory deficits and rescuing the lost of D_1_R-H_3_R complexes at late stages of HD.

## Results

### Functional D_1_R-H_3_R heteromers are expressed in wild type STHdh^Q7^ and HD STHdh^Q111^STHdhstriatal cell model

To test whether D_1_R-H_3_R heteromers could indeed be targets for controlling D_1_R signaling in HD, we first analyzed the expression of both receptors in immortalized striatal cells expressing endogenous levels of full-length wild-type STHdh^Q7^ or mutant STHdh^Q111^ huntingtin (28). Ligand binding determined that both STHdh^Q7^ and STHdh^Q111^ cells endogenously express similar levels of D_1_R and H_3_R (Supplemental Table 1). By proximity ligation assays (PLA), D_1_R-H_3_R heteromers were detected as red spots surrounding the blue stained nuclei in both cell types (Fig. 1A, **left panels of both cell types**) and in cells treated with control lentivirus vector (Fig. S1A) but not in cells depleted of H_3_R (Fig. 1A, **right panels of both cell types**) by shRNA, as shown by RT-PCR and functionality (Fig. S1 B, C), or in negative controls (Fig. S1D). To ensure that D_1_R-H_3_R heteromers were functional in STHdh cells, cell signaling experiments were performed. Using both STHdh^Q7^ and STHdh^Q111^ cells and concentrations of ligands previously shown to be optimal for receptor activation of the ERK1/2 pathway (26;29;30), we observed that the D_1_R agonist SKF 81297 was able to increase ERK1/2 phosphorylation whereas it was prevented by D_1_R antagonist SCH 23390, and by the H_3_R antagonist thioperamide (Fig. S2A, B) via cross-antagonism. In addition, we tested a previously described alternative signaling pathway activated downstream of D_1_R, Ca^2+^ mobilization (31;32). When cells were treated with the D_1_R agonist SKF 81297 a robust and rapid increase in cytosolic Ca^2+^ was detected in both STHdh^Q7^ and STHdh^Q111^ cells (Fig. 1B, C). Importantly, this calcium release could be dampened with the H_3_R antagonist thioperamide (cross-antagonism) (Fig. 1B, C). The above signaling data strongly support the presence of functional D_1_R-H_3_R heteromers in STHdh cells.

**Figure 1.**
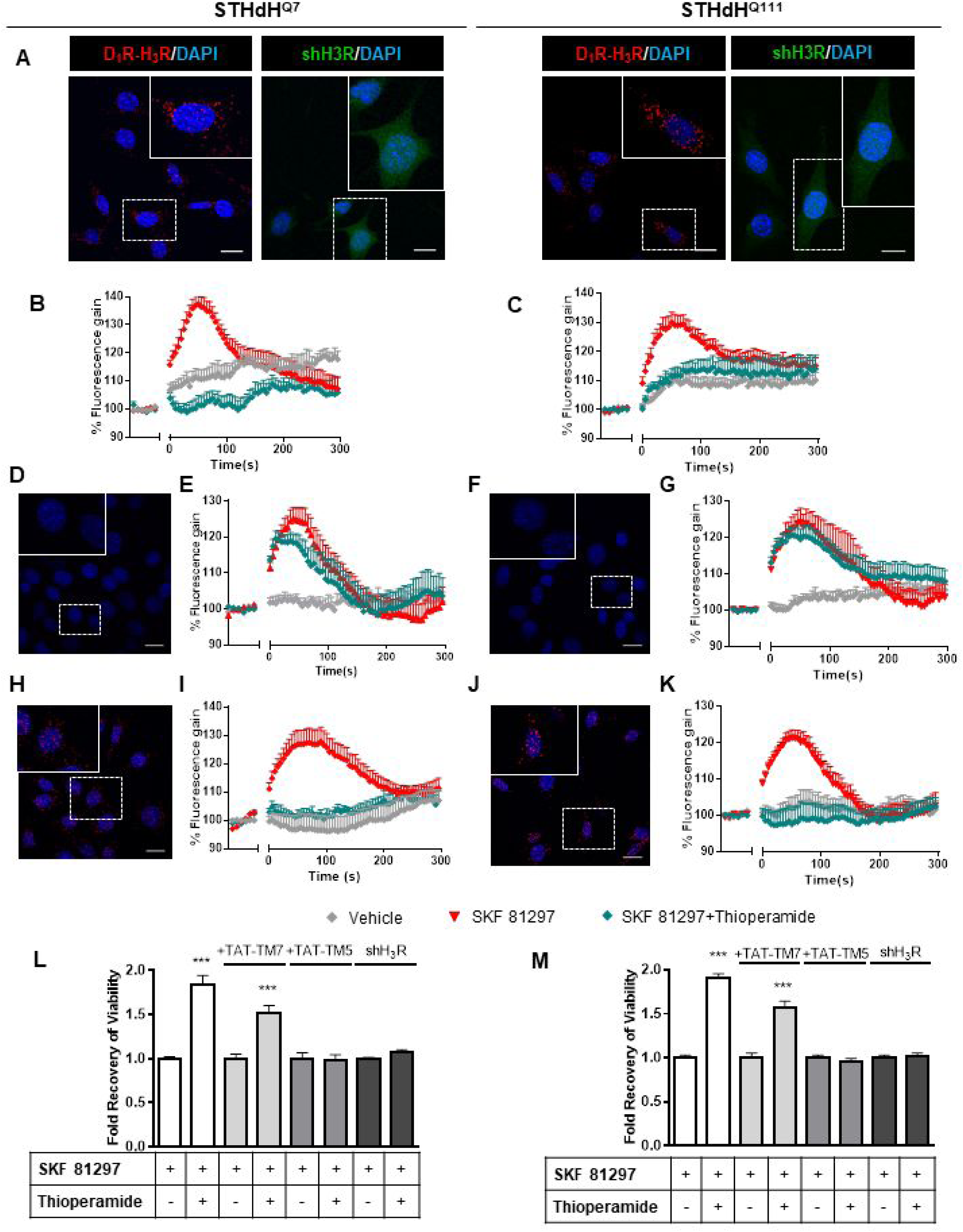
Functional D_1_R-H_3_R heteromers are expressed in STHdH^Q7^ and STHdH^Q111^ cells. PLA were performed in STHdH^Q7^ and STHdH^Q111^ cells (**A, D, F, H and J**) or in cells infected with shH_3_R to silence H_3_R, observed as green stained cells due to the GFP expression included in the plasmid (**A**). D_1_R-H_3_R heteromers were visualized in STHdH cells as red spots around blue colored DAPI stained nucleus, but not in STHdH cells infected with shH_3_R vector (**A**). Calcium increases were measured in STHdH^Q7^ (**B, E and I**) or STHdH^Q111^ (**C, G and K**). Cells were treated (20 min) or not with the H_3_R antagonist thioperamide (10 μM) before the addition of vehicle or SKF 81297 (1 μM). In (**D, E, F, G, H, I, J and K**), STHdH^Q7^ (**D, E, H and I**) or STHdH^Q111^ (**F, G, J and K**) cells were also pre-treated for 60 min with 4 μM TM5 (**D, E, F and G**) or TM7 (**H, I, J and K**) peptides. Heteromers were visualized as red spots around DAPI (blue) stained nucleus in cells pre-treated with TM7 peptide. Scale: 20 μm. For each calcium curve values are expressed as a percentage increase with respect to untreated cells and are a mean ± SEM of 3 to 5 independent experiments. In (**L** and **M**), cell viability was determined in STHdH^Q7^ (**L**) or STHdH^Q111^ cells (**M**) pre-treated for 60 min with vehicle (white columns), with 4 μM TAT-TM7 (pale grey columns) or TAT-TM5 (grey columns) or infected with shH_3_R to silence H_3_R (dark grey columns) prior overstimulation with 30 μM SKF 81297. Values represent mean ± SEM (n = 24 to 30) of cell viability recovery expressed as in-fold respect to SKF 81297 treated cells. Student’s t test showed a significant (***p < 0.001) effect over SKF 81297 treated cells.

To further demonstrate that an H_3_R antagonist is dampening D_1_R activation involving D_1_R-H_3_R heteromers, we evaluated the effect of interfering peptides, which are synthetic peptides with the amino acid sequence of domains of the receptors involved in the heteromeric interface. This approach has been used by us and others to disrupt other heteromer complexes (33–37). In a previous study we showed the efficacy of this approach in demonstrating heteromerization of D_1_R with D_3_R, using a peptide with the sequence of D_1_R transmembrane domain 5 (TM5) but not TM7 (34). We therefore investigated whether synthetic peptides with the sequence of TM5, and TM7 (as a negative control) of D_1_R, fused to HIV-TAT, were also able to disrupt receptor D_1_R-H_3_R heteromers measured by PLA. In agreement with our hypothesis, there was a near complete loss in PLA fluorescence signal when STHdh^Q7^ and STHdh^Q111^ cells were incubated with TAT-TM 5 peptide (Fig. 1D, F), but not for the negative control in which the TAT-TM 7 peptide was used (Fig. 1H, J). We next evaluated whether TM5 or TM7 would interfere with the observed cross-antagonism in calcium mobilization assays. Clearly, pretreatment of both STHdh^Q7^ and STHdh^Q111^ cells with the TAT-TM5 (Fig. 1E, G) but not TAT-TM7 (Fig. 1I, K) peptide disrupts the ability of the H_3_R antagonist thioperamide to dampen D_1_R calcium signaling. These results support that TM5 forms part of the interface of the D_1_R-H_3_R heteromer and demonstrate that the H_3_R antagonist effect is driven through direct interaction between D_1_R and H_3_R.

### H_3_R ligands prevent the D_1_R-induced cell death in STHdh^Q7^ and STHd^Q111^ cells

It has been previously reported that upon activation of D_1_R, STHdh cell viability is reduced (10). To explore whether H_3_R ligands could impair D_1_R activation through D_1_R-H_3_R heteromers in a pathologically relevant readout, we used D_1_R-induced cell death as an output of D_1_R activation in STHdh cells. As expected, STHdh cell viability decreased when treated with the D_1_R agonist SKF 81297 in a concentration-dependent manner (Fig. S2C). Significant cell death did not occur until 30 μM SKF 81297 was used (Fig.S2C), an effect prevented by the D_1_R antagonist SCH 23390 (Fig. S2E). Pre-treatment with the H_3_R antagonist thioperamide, which did not modify cell viability when administered alone (Fig. S2E), increased the number of surviving cells in the presence of the D_1_R agonist SKF 81297 in both cell types (Fig. 1L, M **and** Fig. S2D). Importantly, the effect of the H_3_R antagonist thioperamide was specific since no protection from D_1_R agonist-induced cell death was observed in cells depleted of H_3_R with shRNA lentiviral infection (Fig. 1L, M), but was observed in cells transfected with the control lentivirus (Fig. S2F). In addition, we also demonstrated that recovery of viability induced by the H_3_R antagonist thioperamide was mediated by D_1_R-H_3_R heteromers since pre-incubation with D_1_R TM5 peptide, but not D_1_R TM7 impaired the H_3_R antagonist protection from D_1_R agonist-induced cell death (Fig. 1L, M).

To better understand the mechanisms involved in D_1_R-H_3_R heteromer action, we determined which cellular signaling pathways are implicated in the cross-antagonism of H_3_R upon activation of D_1_R. Both concentrations of the D_1_R agonist SKF 81297, cytotoxic (30 µM) and non-cytotoxic (1 µM), can induce intracellular calcium release, which is more pronounced and persistent at 30 μM (Fig. S4A and B). As occurred at 1 μM SKF 81297 (see Fig. 1), the calcium release induced by 30 μM SKF 81297 was also blocked by the H_3_R antagonist thioperamide (Fig. S3A and B). A correlation between the intensity of calcium responses and the activation of apoptotic pathways such as p38 (38) has been previously demonstrated. Thus, we measured changes in p38 phosphorylation levels using both concentrations of the D_1_R agonist SKF 81297 (Fig. S4C and D). Interestingly, we found that increased phosphorylation of p38 only occurred at the cytotoxic concentration of SKF 81297. Treatment with the H_3_R antagonist thioperamide reduced p38 phosphorylation upon D_1_R activation in both cell types (Fig. S3C). Moreover, the p38 inhibitor SB 203580 blocked p38 phosphorylation (Fig. S3C) and protected against the cytotoxic effect of the D_1_R agonist SKF 81297 in a dose-dependent manner (Fig. S3D), confirming that p38 is a key pathway involved in D_1_R-mediated cell death in these cells.

Overstimulation of D_1_R induces receptor internalization promoting rapid intracellular signaling (39), while D_1_R expression is decreased in several models of HD (40). Receptor internalization can activate secondary signaling pathways (41). To test whether changes in receptor trafficking might be at play we analyzed whether 30 μM SKF 81297 can induce D_1_R internalization in the striatal cells. We observed that 30 μM SKF 81297, that decreased cell viability, promoted D_1_R internalization in both STHdh cells (Fig. S5A). Interestingly, the 30 μM SKF 81297-induced D_1_R internalization correlated with D_1_R-H_3_R heteromer disruption evidenced by a lack of PLA staining in both STHdh cells treated with 30 μM SKF 81297 (Fig. S5B). One potential way by which GPCRs can influence each other in a heteromer is by altering the trafficking of the partner receptor (42). Pre-treatment with the H_3_R antagonist thioperamide restored the number of punctate PLA spots decreased after overstimulation with the D_1_R agonist SKF 81297 (Fig. S5B). These results suggest that H_3_R ligands are impede D_1_R internalization and D_1_R-mediated cell death by inhibiting p38 phosphorylation and calcium signaling.

### Functional D_1_R-H_3_R heteromers are expressed in wild-type Hdh^Q7/Q7^ and in Hdh^Q7/Q111^ mutant knock-in mice at early but not late HD stages

To test whether D_1_R-H_3_R heteromers can indeed be targets for treating HD, we investigated their expression and function in the striatum, cerebral cortex and hippocampus of a widely accepted preclinical model of HD, the heterozygous Hdh^Q7/Q111^ mutant knock-in mice, and their wild-type Hdh^Q7/Q7^ littermates (43;44). By PLA we confirmed that both Hdh^Q7/Q7^ and Hdh^Q7/Q111^ mice display D_1_R-H_3_R heteromers at 2 months (mo) (Fig. S6) and 4 mo (Fig. 2A) of age in all brain regions tested. No signal was observed in negative controls in which one of the PLA primary antibodies were missing (Fig. S7). Heteromer expression was similar in all brain areas and no differences were observed between genotypes at 4 mo of age (Fig. 2B). Surprisingly, an almost complete loss of D_1_R-H_3_R heteromers was found in 6 mo and 8 mo-old Hdh^Q7/Q111^ mice but not in Hdh^Q7/Q7^ mice (Fig. S8 **and** Fig. 3A and B), indicating that at more advanced disease stages the D_1_R-H_3_R heteromer is lost. Although at 8 mo of age we detected a partial decrease in striatal D_1_R expression in Hdh^Q7/Q111^ compared with Hdh^Q7/Q7^ mice using ligand binding experiments **(Supplemental Table 2)**, the loss of heteromer expression is not due to a complete loss of receptor expression since by radioligand binding **(Supplemental Table 2)** and mRNA expression analysis **(Supplemental Table 3)** both receptors continue to be expressed.

**Figure 2.**
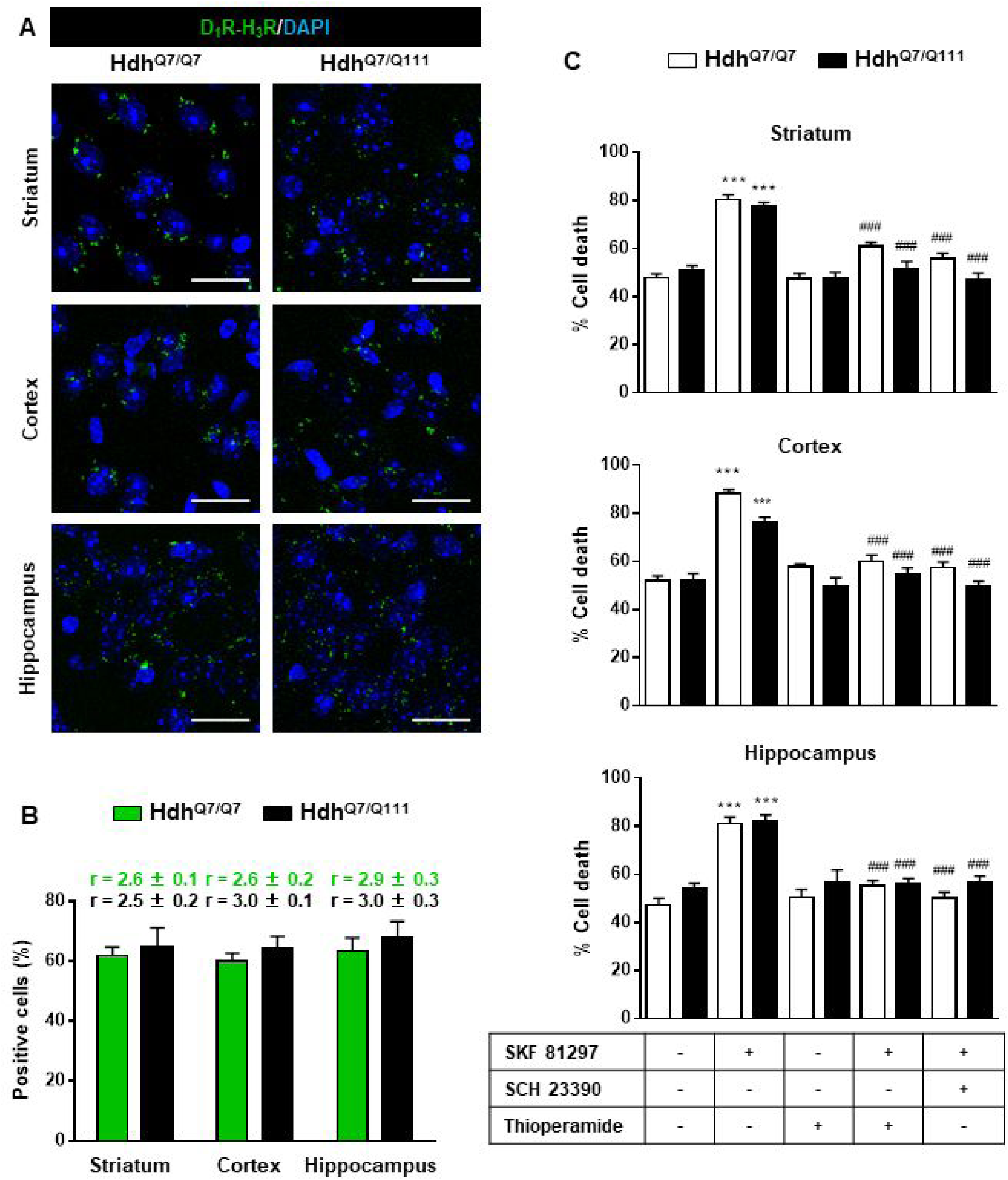
Functional D_1_R-H_3_R heteromers are expressed in wild-type Hdh^Q7/Q7^ and mutant Hdh^Q7/Q111^ mice. Striatal, cortical or hippocampal slices from 4-month-old Hdh^Q7/Q7^ and Hdh^Q7/Q111^ mice were used. In (**A**), by Proximity Ligation Assays (PLA) D_1_R-H_3_R heteromers were visualized in all slices as green spots around blue colored DAPI stained nucleus. Scale bar: 20 μm. In (**B**), the number of cells containing one or more green spots is expressed as the percentage of the total number of cells (blue nucleus). *r values* (number of green spots/cell containing spots) are shown above each bar. Data (% of positive cells or r) are the mean ± SEM of counts in 600-800 cells from 4-8 different fields from 3 different animals. Student’s t test showed no significant differences in heteromers expression in Hdh^Q7/Q7^ and Hdh^Q7/Q111^ mice. In (**C**), striatal, cortical or hippocampal organotypic slice cultures from 4-month-old Hdh^Q7/Q7^ and Hdh^Q7/Q111^ mice were treated for 60 min with vehicle, the D_1_R antagonist SCH 23390 (10 μM) or H_3_R antagonist thioperamide (10 μM) before the addition of SKF 81297 (50 μM). After 48h cell death was determined. Values represent mean ± SEM (n = 3 to 19) of percentage of cell death. One-way ANOVA followed by Bonferroni post hoc tests showed a significant effect over non-treated organotypic cultures (***p < 0.001) or of the H_3_R antagonist plus SKF 81297 treatment over the SKF 81297 (^###^p < 0.001).

**Figure 3.**
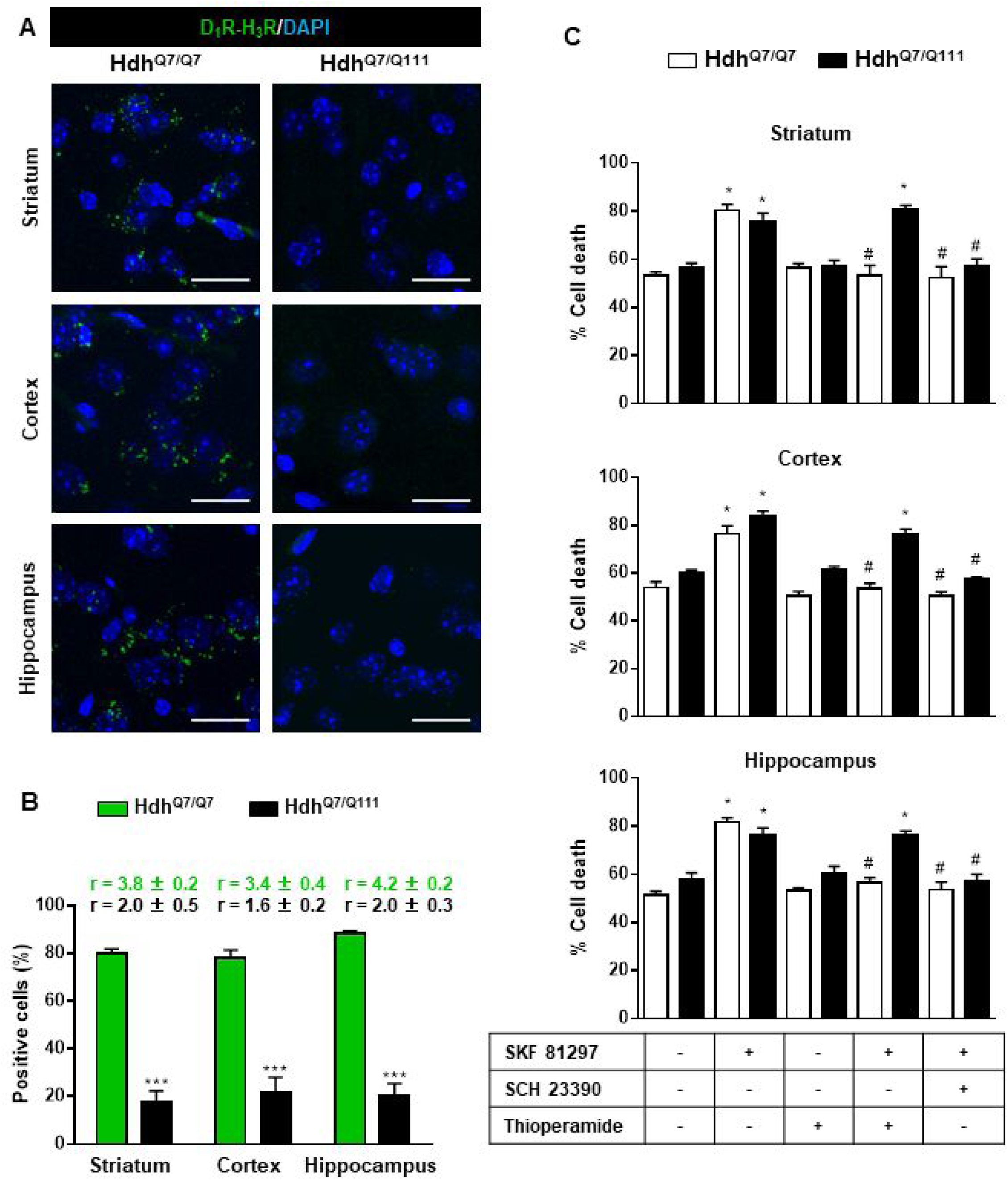
Functional D_1_R-H_3_R heteromers are expressed in wild-type Hdh^Q7/Q7^ but not in 8-month-old mutant Hdh^Q7/Q111^ mice. Striatal, cortical or hippocampal slices from 8-month-old Hdh^Q7/Q7^ and Hdh^Q7/Q111^ mice were used. In (**A**), by Proximity Ligation Assays (PLA) D_1_R-H_3_R heteromers were visualized in Hdh^Q7/Q7^ mice but not in Hdh^Q7/Q111^ mice as green spots around blue colored DAPI stained nucleus. Scale bar: 20 μm. In (**B**), the number of cells containing one or more green spots is expressed as the percentage of the total number of cells (blue nucleus). *r values* (number of green spots/cell containing spots) are shown above each bar. Data (% of positive cells or r) are the mean ± SEM of counts in 600-800 cells from 5-7 different fields from 3 different animals. Student’s t test showed a significant (***p<0.05) decrease of heteromers expression in Hdh^Q7/Q111^ mice compared to the respective Hdh^Q7/Q7^ mice. In (**C**) striatal, cortical or hippocampal organotypic slice cultures from 8-month-old Hdh^Q7/Q7^ and Hdh^Q7/Q111^ mice were treated for 60 min with medium, the D_1_R antagonist SCH 23390 (10 μM) or the H_3_R antagonist thioperamide (10 μM) before the addition of SKF 81297 (50 μM) and cell death was determined. Values represent mean ± SEM (n = 3 to 6) of percentage of cell death. One-way ANOVA followed by Bonferroni post hoc tests showed a significant effect over non-treated organotypic cultures (*p < 0.05) or of the H_3_R antagonist plus SKF 81297 treatment over the SKF 81297 (^#^p < 0.05).

To test the role of D_1_R-H_3_R heteromers, organotypic mouse striatal, cortical and hippocampal cultures were obtained. Cell death was induced by the D_1_R agonist SKF 81297 (50 µM), and analysis of DAPI and propidium iodide staining was performed. As expected, D_1_R agonist SKF 81297 treatment increased the percentage of cell death in all three regions compared to vehicle-treated organotypic cultures without significant differences between genotypes at 4 mo of age (Fig. 2C). Importantly, slices pre-treated with the H_3_R antagonist thioperamide, that does not modify cell death when administered alone, protected cells from D_1_R elicited cell death (Fig. 2C), indicating that functional D_1_R-H_3_R heteromers are expressed in different brain areas of Hdh^Q7/Q7^ and Hdh^Q7/Q111^ mice at early disease stages. The dramatic change in heteromer expression in 8 mo-old Hdh^Q7/Q111^ mice was mirrored by the lack of protection of the H_3_R antagonist thioperamide against SKF 81297-induced cell death in organotypic cultures (Fig. 3C), corroborating that the presence of D_1_R-H_3_R heteromers is needed for the H_3_R antagonist to prevent D_1_R-mediated cell death.

### Treatment with thioperamide prevents cognitive and motor learning deficits at early disease stages

To test whether the H_3_R antagonist thioperamide can exert beneficial effects in the initial stages of the disease we evaluated the effect of chronic thioperamide treatment on motor learning and memory deficits in mutant Hdh^Q7/Q111^ mice. Since cognitive decline is observed in these HD mice from 6 mo of age (43–45) and the D_1_R-H_3_R heteromers are expressed and functional until the age of 5 mo (Fig. S9A-D), we chose 5 mo-old animals to start the thioperamide treatment (Fig. S10). Corticostriatal function in saline and thioperamide-treated Hdh^Q7/Q7^ and Hdh^Q7/Q111^ mice was analyzed by using the accelerating rotarod task that evaluates the acquisition of new motor skills (44). Saline-treated mutant Hdh^Q7/Q111^ mice were unable to maintain their balance on the rotarod as wild-type Hdh^Q7/Q7^ mice revealing impaired acquisition of new motor skills (Fig. 4A). Chronic treatment with thioperamide completely rescued motor learning deficits in mutant Hdh^Q7/Q111^ mice as evidenced by a similar latency to fall in the accelerating rotarod as wild type Hdh^Q7/Q7^ mice. Next, recognition long-term memory (LTM) was analyzed by using the novel object recognition test (NORT) (Fig. 4B). After two days of habituation in the open field arena (Fig. S11A, B, C, D **and** Fig. S10E, F, G, H), no significant differences were found between genotypes and/or treatments, demonstrating no alterations in motivation, anxiety or spontaneous locomotor activity. After habituation, animals were subjected to a training session in the open field arena in the presence of two similar objects (A and A’). Both saline and thioperamide-treated wild-type Hdh^Q7/Q7^ and mutant Hdh^Q7/Q111^ mice similarly explored both objects indicating neither object nor place preferences (Fig. 4B). After 24 h, LTM was evaluated by changing one of the old objects (A’) for a novel one (B). Whereas saline-treated Hdh^Q7/Q111^ mice did not show any preference for the novel object with respect to the familiar one, indicating recognition LTM deficits, thioperamide treatment completely prevented this LTM deficit in mutant Hdh^Q7/Q111^ mice (Fig. 4B). Next, spatial LTM was analyzed using the T-maze spontaneous alternation task (T-SAT) (Fig. 4C). During the training, similar exploration time (Fig. 4C, **left panel)** and similar number of arm entries (Fig. S12, **left panel)** were found in all genotypes and treatments. After 5 h, a testing session showed that saline-treated Hdh^Q7/Q111^ mice had no preferences between the novel arm and the old arm, indicating spatial LTM deficits (Fig. 4C, **right panel)**. Interestingly, mutant Hdh^Q7/Q111^ mice treated with thioperamide spent more time in the novel *versus* the old arm, revealing preserved LTM (Fig. 4C, **right panel)**. Overall, these data demonstrate the effectiveness of thioperamide treatment in restoring motor learning and preventing spatial and recognition LTM deficits in mutant Hdh^Q7/Q111^ mice.

**Figure 4.**
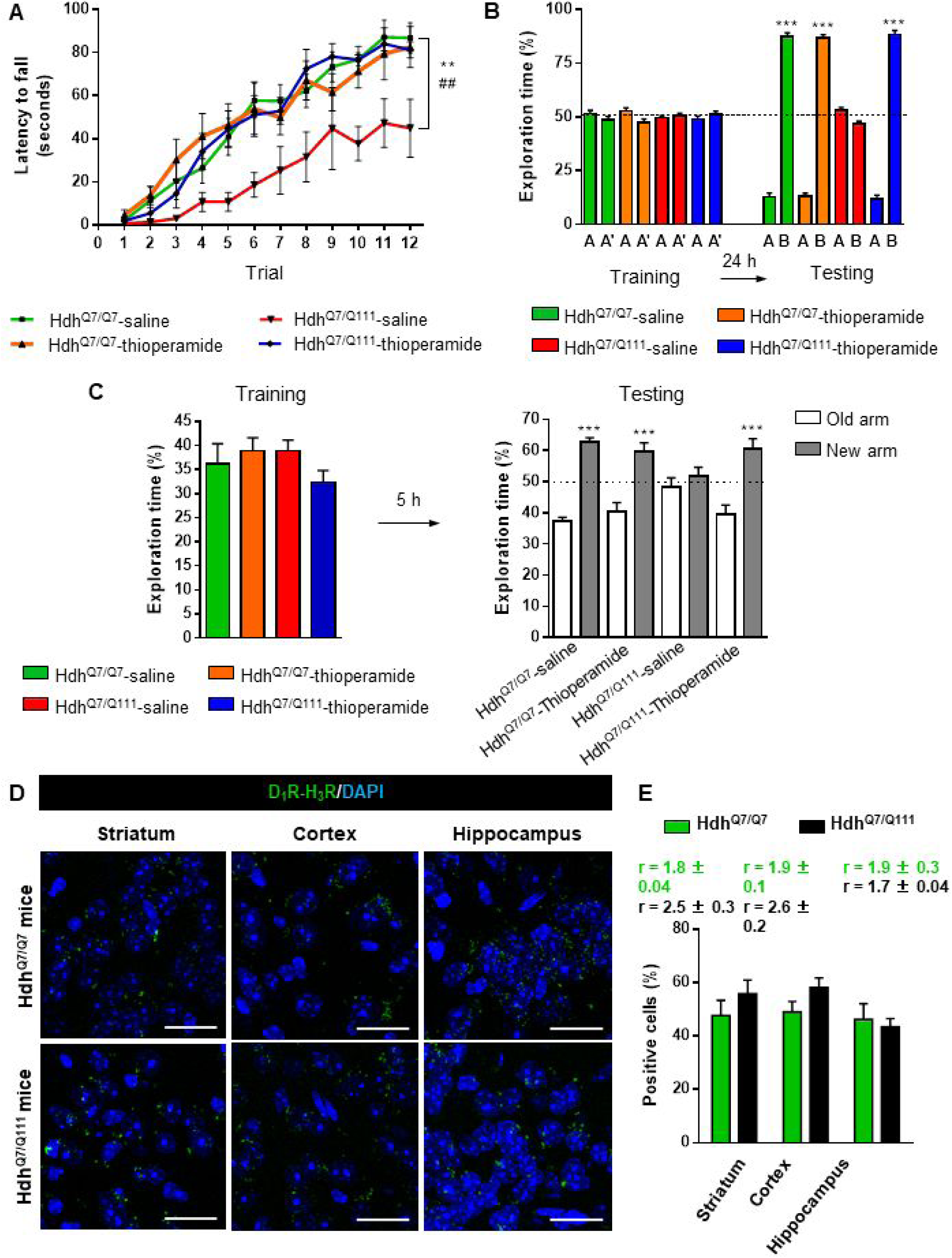
Thioperamide chronic treatment prevents motor learning, long-term memory (LTM) deficits and the loss of receptor heteromerization in 6-month-old Hdh^Q7/Q111^ mice. In (**A**), curves illustrating the latency to fall in the accelerating rotarod of 6-month-old Hdh^Q7/Q7^ and Hdh^Q7/Q111^ mice treated with saline or thioperamide from 5 months of age are shown. In (**B**), the exploration time for saline or thioperamide-treated Hdh^Q7/Q7^ and Hdh^Q7/Q111^ mice during the training and the testing (24 h delay, LTM) sessions in a novel-object recognition task showing that long-term recognition memory deficits are rescued in the thioperamide-treated Hdh^Q7/Q111^ mice. One-way ANOVA with Bonferroni *post hoc* showed significant differences (***p<0.001) compared to the old object recognition. In (**C**), bar diagram illustrating the exploration time for saline- or thioperamide-treated Hdh^Q7/Q7^ and Hdh^Q7/Q111^ mice during the training and the 5 h later testing in the T-SAT showing thioperamide reverses spatial long-term memory (LTM) deficits. In (**a** to **c**), 11 saline-treated Hdh^Q7/Q7^ mice, 10 thioperamide-treated Hdh^Q7/Q7^ mice, 7 saline-treated Hdh^Q7/Q111^ mice and 9 thioperamide-treated Hdh^Q7/Q111^ mice were evaluated at 6 months of age. In (**D**) PLA were performed in striatal, cortical and hippocampal slices from 6-month-old Hdh^Q7/Q7^ and Hdh^Q7/Q111^ mice treated with thioperamide. D_1_R-H_3_R heteromers were visualized in all samples as green spots around blue colored DAPI stained nucleus. Scale bar: 20 μm. In (**E**) the right panel, the number of cells containing one or more green spots is expressed as the percentage of the total number of cells (blue nucleus). *r values* (number of green spots/cell containing spots) are shown above each bar. Data (% of positive cells or r) are the mean ± SEM of counts in 600-800 cells from 4-8 different fields from 3 different animals. Student’s *t* test showed no significant differences in heteromer expression in thioperamide-treated Hdh^Q7/Q111^ mice compared to the respective Hdh^Q7/Q7^ mice.

We next tested if the reversion of the HD phenotype in mutant Hdh^Q7/Q111^ mice induced by thioperamide treatment correlated with the preservation of D_1_R-H_3_R heteromer expression. By PLA we observed that in saline-treated 6-mo-old Hdh^Q7/Q111^ mice the heteromer expression was significantly diminished with respect to the age-matched Hdh^Q7/Q7^ mice (Fig. S8A and B). Notably, treatment with thioperamide significantly prevented the loss of D_1_R-H_3_R heteromers in all brain regions analyzed in Hdh^Q7/Q111^ mice at both 6 (Fig. 4D and E) and 8 mo of age (Fig. S13A and B), suggesting that the altered trafficking observed in cells may potentially also occur in vivo.

### Treatment with thioperamide ameliorates spinophilin-immunoreactive puncta alterations in the motor cortex and hippocampus of 6-month-old mutant Hdh^Q7/Q111^ _mice_

Alterations in dendritic spine dynamics, density and morphology are critically involved in the synaptic deficits present in HD (4;44-51). We recently described a significant decrease in dendritic spine density in the hippocampus (45) and the motor cortex of mutant Hdh^Q7/Q111^ mice (44) without significant alterations in the striatum. To analyze whether the improvement of motor learning and memory deficits observed in thioperamide-treated mutant Hdh^Q7/Q111^ mice was associated with a recovery in the density of dendritic spines, spinophilin immunostaining was performed in CA1 hippocampal and motor cortical coronal slices obtained from 6-mo-old wild-type Hdh^Q7/Q7^ and mutant Hdh^Q7/Q111^ mice (Fig. 5A and B **and** Fig. S14A). This methodology was used by us and others to identify structural alterations in dendritic spines (44;52;53). Confocal microscopy analyses revealed a significant reduction in the density of spinophilin-immunoreactive puncta in the *stratum radiatum* (apical dendrites of CA1 pyramidal neurons) and *stratum oriens* (basal dendrites of CA1 pyramidal neurons) of saline-treated 6-mo-old mutant Hdh^Q7/Q111^ mice compared to saline-treated wild-type Hdh^Q7/Q7^ mice (Fig. 5A **and** Fig. S14A). Interestingly, thioperamide treatment prevented the decline in the number of spinophilin-immunoreactive puncta in mutant Hdh^Q7/Q111^ mice. Similar data was obtained when the layers of the motor cerebral cortex (M1) were analyzed. A significant reduction in the density of spinophilin-immunoreactive puncta in layer I and layer II-III, but not layer V, of the motor cortex of 6-mo-old saline-treated Hdh^Q7/Q111^ mice was found compared to saline-treated Hdh^Q7/Q7^ mice (Fig. 5B **and** Fig. S14A). Interestingly, thioperamide-treated Hdh^Q7/Q111^ mice exhibited a complete recovery in the density of spinophilin-immunoreactive puncta (Fig. 5A, 5B **and** Fig. S14A). No significant differences were found between groups when the mean size of spinophilin puncta was analyzed (Fig. S14A). Altogether, these data demonstrate that the loss of spinophilin immunoreactive-puncta in mutant Hdh^Q7/Q111^ mice can be ameliorated by thioperamide treatment.

**Figure 5.**
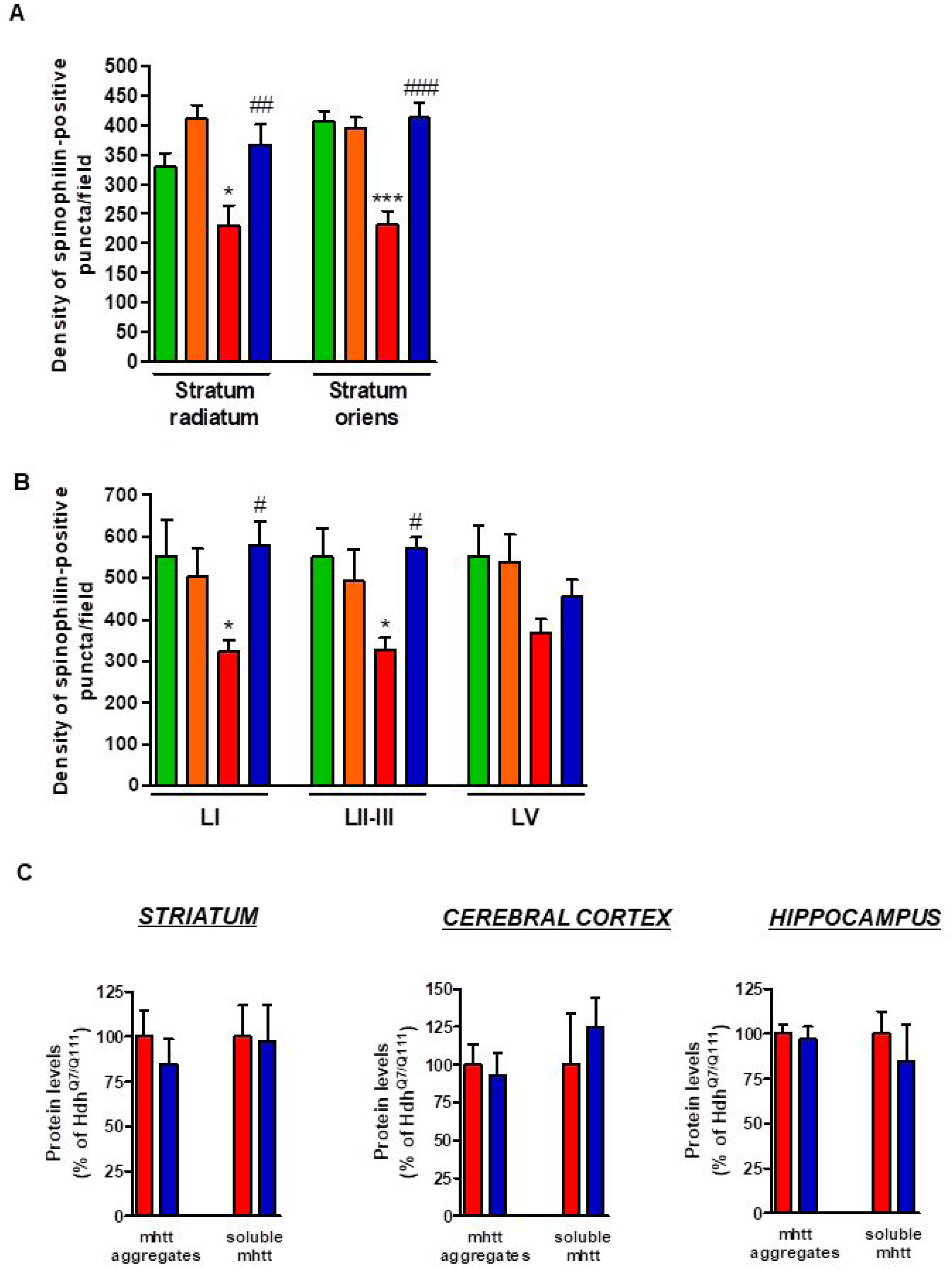
Thioperamide treatment restored spinophilin-immunoreactive puncta reduction in the hippocampus and motor cortex of Hdh^Q7/Q111^ mice and exerts no effect on the clearance of mutant huntingtin accumulation. In (**A**) spinophilin-immunoreactive puncta were counted in the *stratum oriens* and *stratum radiatum* of CA1 hippocampus and in (**B**) layers I, II/III and V of motor cortex area 1 (M1) of saline and thioperamide-treated WT HdhQ7/Q7 and knock-in HdhQ7/Q111 mice. Quantitative analysis is shown as mean ± SEM (n= 9 images from three animals/group). Statistical analysis was performed using Student’s two-tailed t test. *p<0.05, ***p<0.001 compared to saline-treated Hdh^Q7/Q7^ mice. #p<0.05, ##p<0.01, ###p<0.001 compared to saline-treated HdhQ^7/Q111^ mice. In (**C**), Quantification of the protein levels of insoluble mHtt oligomeric forms and soluble mHtt forms of total striatal, hippocampal and cortical extracts from 6-month-old saline and thioperamide-treated knock-in Hdh^Q7/Q111^ mice analysed by immunoblot. All histograms represent the mean ± SEM (n=6-8 per group). Student’s *t* test showed no significant differences between groups.

We also evaluated mutant huntingtin (mhtt) aggregates in the striatum, cerebral cortex and hippocampus of mutant Hdh^Q7/Q111^ mice after saline or thioperamide treatment, as another pathological hallmark of HD (43;54;55). 1C2 immunostaining revealed in lysates from either vehicle ortreated mutant Hdh^Q7/Q111^ mice a substantial accumulation of mhtt oligomeric forms detected as a diffuse smear in the stacking gel (Fig. S14B). Thioperamide treatment failed to prevent the accumulation of these oligomeric forms (Fig. 5C **and** Fig. S14B). No significant differences between groups were found when soluble monomeric mhtt levels were analyzed (Fig. 5C **and** Fig. S14B).

### Thioperamide treatment does not rescue memory and motor learning deficits in mutant Hdh^Q7/Q111^ mice when D_1_R-H_3_R heteromers are lost

If the behavioral improvements observed after thioperamide treatment are mediated by the D_1_R-H_3_R heteromer and not just by the blockade of the single H_3_R, then a treatment paradigm in the absence of the heteromer should have no effect. To test this hypothesis, we used wild-type Hdh^Q7/Q7^ and mutant Hdh^Q7/Q111^ mice at the age of 7 months, when we found the heteromer to be lost. Animals were chronically treated with saline or thioperamide for 1 month and motor learning was evaluated using the accelerating rotarod task. As expected, saline-Hdh^Q7/Q111^ mice exhibited poor performance in this task showing shorter latency to fall compared to wild-type Hdh^Q7/Q7^ mice (Fig. 6A). Notably, thioperamide treatment had no effect on motor learning performance as both saline- and thioperamide-treated mutant Hdh^Q7/Q111^ mice were indistinguishable demonstrated by similar latency to fall in the accelerating rotarod task (Fig. 6A).

**Figure 6.**
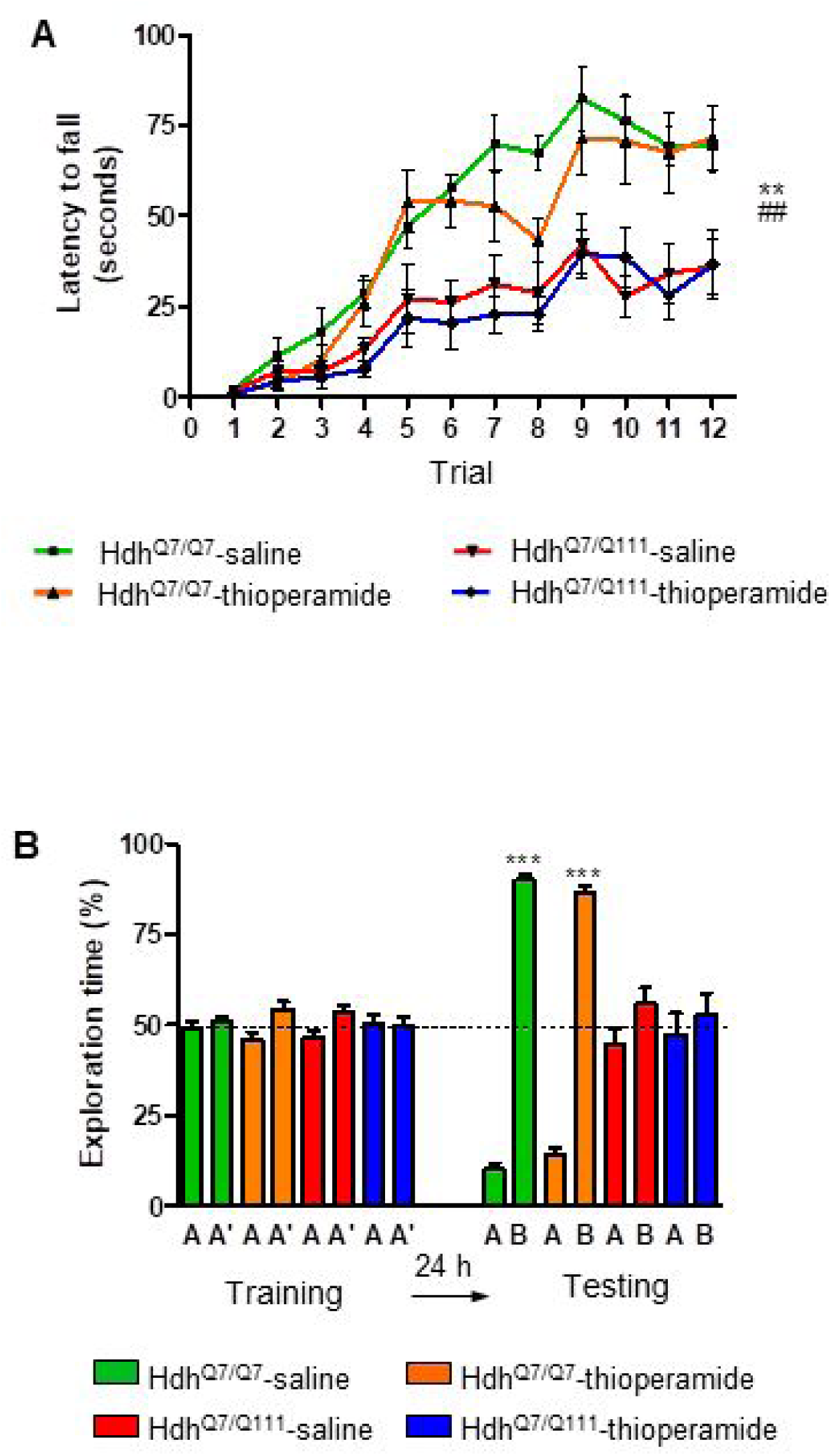
Thioperamide chronic treatment does not prevent motor learning and long-term memory (LTM) deficits in 8-month-old Hdh^Q7/Q111^ mice when the D_1_R-H_3_R heteromer is not expressed. In (**A**), curves illustrating the latency to fall in the accelerating rotarod of 8-month-old Hdh^Q7/Q7^ and Hdh^Q7/Q111^ mice treated with saline or thioperamide from 7 months of age are shown. Two-way ANOVA with repeated measures showed significant differences (**p<0.01) of saline-treated Hdh^Q7/Q111^ mice compared to saline-treated Hdh^Q7/Q7^ mice or (##p<0.01) thioperamide-treated Hdh^Q7/Q111^ mice compared to saline-treated Hdh^Q7/Q7^ mice. 11 saline-treated Hdh^Q7/Q7^ mice, 11 thioperamide-treated Hdh^Q7/Q7^ mice, 8 saline-treated Hdh^Q7/Q111^ mice and 9 thioperamide-treated Hdh^Q7/Q111^ mice were evaluated at 8 months of age. In (**B**), bar diagram illustrating the exploration time for saline or thioperamide-treated Hdh^Q7/Q7^ and Hdh^Q7/Q111^ mice during the training and the testing (24 h delay, LTM) sessions in a novel-object recognition task showing that long-term recognition memory deficits are not rescued in the thioperamide-treated Hdh^Q7/Q111^ mice. One-way ANOVA with Bonferroni *post hoc* comparisons showed significant differences (***p<0.001) compared to the old object recognition. 11 saline-treated Hdh^Q7/Q7^ mice, 12 thioperamide-treated Hdh^Q7/Q7^ mice, 10 saline-treated Hdh^Q7/Q111^ mice and 11 thioperamide-treated Hdh^Q7/Q111^ mice were evaluated at 8 months of age.

We next asked whether thioperamide treatment could improve cognitive function by rescuing memory deficits in these same animals. Saline-treated 8-mo-old Hdh^Q7/Q111^ mice exhibited long-term memory deficits when recognition memory was analyzed using the novel object recognition test (NORT) (Fig. 6B). Similar to motor learning results, chronic treatment with thioperamide did not rescue Hdh^Q7/Q111^ mice from memory deficits (Fig. 6B). Overall, these results demonstrate that the effect of thioperamide in learning and memory in Hdh^Q7/Q111^ mice requires the proper expression and function of D_1_R-H_3_R heteromers.

### D_1_R-H_3_R heteromer expression changes occur in other rodent HD models and in HD patients

The fact that thioperamide treatment 1) prevents cognitive and motor learning deficits, 2) ameliorates striatal neuropathology, 3) ameliorates morphological alterations and 4) prevents the loss of D_1_R-H_3_R heteromers at 6 mo and 8 mo of age in a mouse model of HD is suggestive that thioperamide, or a future pharmacologically improved H_3_R antagonist specifically targeting D_1_R-H_3_R heteromers, can be used to treat HD symptoms. To test this, we investigated D_1_R-H_3_R heteromer expression in other transgenic HD mouse models and in human caudate-putamen slices using PLA. The loss of heteromer expression compared with wild-type littermates was also observed in other mouse models of HD, the R6/1 and R6/2 mice transgenic for the human huntingtin exon 1 (Fig. S15A and B, **respectively)**. Importantly, D_1_R-H_3_R heteromers were detected as green spots surrounding the blue stained nuclei in human caudate-putamen slices from control individuals and low-grade (grade 0, 1 and 2) HD patients (Fig. 7A and B). In contrast, green spots were almost absent in samples from high-grade (grade 3 or grade 4) HD patients (Fig. 7A and B). These results show that D_1_R-H_3_R heteromer formation changes during disease progression and, importantly, that humans express D_1_R-H_3_R heteromers at early disease stages.

**Figure 7.**
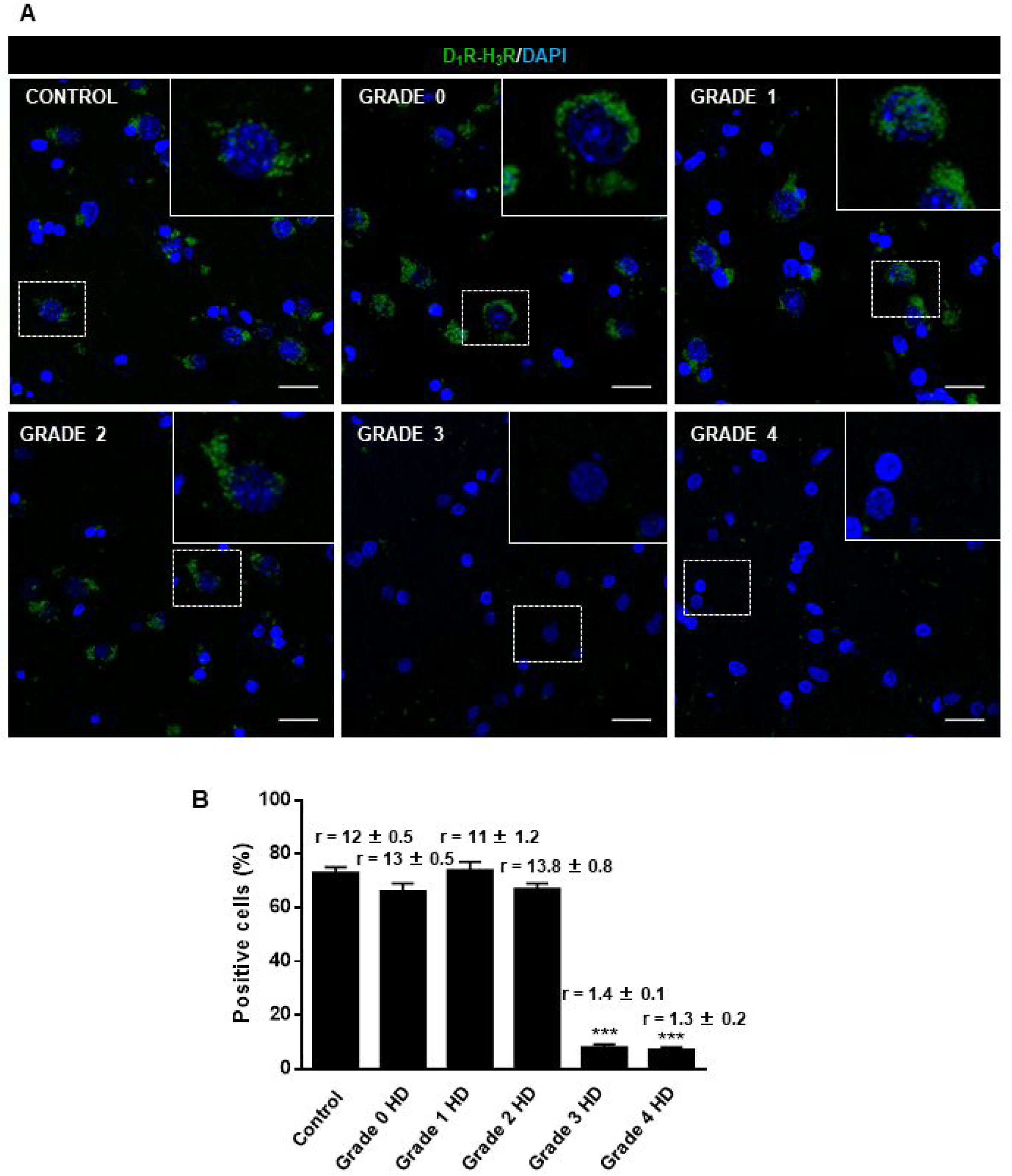
Striatal D_1_R-H_3_R heteromers are expressed in human control subjects and grade 2 HD patients but not in grade 3-4 HD patients. In (**A**), by Proximity Ligation Assays (PLA), D_1_R-H_3_R heteromers were visualized as green spots around blue colored DAPI stained nucleus in human striatal slices from age matched control subjects and 0-2 grade HD patients but not in 3-4 grade HD patients. Scale bar: 20 μm. In (**B**), the number of cells containing one or more green spots is expressed as the percentage of the total number of cells (blue nucleus). *r values* (number of green spots/cell containing spots) are shown above each bar. Data are mean ± SEM of counts in 600-800 cells from 10 different fields from subject described in Materials and Methods. Student’s t test showed a significant (***p<0.001) decrease of heteromers expression in 3-4 grade HD patients compared to control subjects.

## Discussion

The imbalance of dopamine inputs throughout HD progression represents a potential “point of no return” for HD patients as this disequilibrium can eventually lead to substantial neuronal dysfunction and cell death. In the present study we demonstrate that 1) excess dopamine signaling via D_1_R leads to cell death by activating the p38 pathway; 2) D_1_R-H_3_R complexes are found within the striatum, cortex and hippocampus of WT mice and in HD mice at early but not late disease stages; 3) targeting D_1_R via D_1_R-H_3_R complexes can slow progression of the disease in early but not late stages when the complexes are lost; and 4) D_1_R-H_3_R complexes are expressed in the human brain and thus represent potential therapeutic targets. This is the first demonstration of GPCR heteromers as potential targets to treat HD. Together, these data support a novel role for D_1_R-H_3_R complexes in neuroprotection and HD.

Several studies have revealed that dopamine neurotoxicity increases the sensitivity of MSSNs to glutamate inputs and leads to striatal neurodegeneration, a role ascribed to aberrant D_1_R and not D_2_R (10;14;56;57). Thus, pharmacological treatments aimed to reduce D_1_R signaling may be beneficial to prevent or slow striatal cell death. Although we cannot rule out the participation of D_2_R in striatal degeneration, our results suggest that D_1_R is a major executor of the final signaling cascades that lead to cell death in HD. This is further supported by the fact that D_1_R is in excess over D_2_R in the striatum, so it is plausible that the former will be more significantly activated than the latter at increased DA levels. We have demonstrated that a toxic but not sub-toxic concentration of SKF81297 increases cytosolic calcium levels and activates the p38 pro-apoptotic pathway. Accordingly, p38 inhibitors completely abrogated the cell death induced by SKF81297 treatment, supporting the benefits of modulation of D_1_R signaling as potential treatment in HD. However, direct manipulation of DA production and/or D_1_R signalling via a specific antagonist has limited therapeutic ability due to associated deleterious side effects. An alternative approach is to modify D_1_R signalling via the histamine neuromodulator. An interaction between H_3_R and the dopaminergic system has been previously reported by us and others (58–60). In this frame, we have demonstrated that H_3_R ligands completely abrogate striatal cell death induced by D_1_R, likely by inhibition of D_1_R-mediated calcium influx and p38 activation. Importantly, D_1_R-H_3_R complexes were found in the striatum, cortex and hippocampus from wild-type Hdh^Q7/Q7^ and mutant Hdh^Q7/Q111^ mice, regions known to be affected by mutant huntingtin toxicity (2;61;62).

The mechanisms of action of D_1_R-H_3_R heteromers can be multiple including allosteric effects. Indeed, the efficacy of the disrupting peptides supports protein-protein-driven effects. A second and potentially additional mechanism is that heteromer formation may alter the trafficking of D_1_R, which could have pleiotropic consequences on signaling. The signaling effects we observe appears to be on a variety of concentrations and timescales in agreement with previous studies showing that GPCR signaling occurs with varied kinetics (63;64). Indeed, part of the concern of trying to target GPCR heteromers for therapeutic purposes is the uncertainty around their stability and thus indirectly whether they can impact GPCR signaling at every timescale. For the case of D_1_R-H_3_R heteromers, it appears that they are stable enough that they can affect both rapid receptor signaling (e.g., Ca^2+^ mobilization) and longer cell signaling pathways like p38, two events that have previously been involved in neuronal cell death in HD (65–69).

Our findings do not rule out that H_3_R ligands by targeting D_2_R-H_3_R heteromers (70) could block D_2_R signaling and contribute to cell death protection. However, several findings argue in favor of D_1_R-H_3_R heteromer as uniquely responsible for the effects of thioperamide on cell death reduction. First, D_1_R over-activation induces cell death-related pathways, D_1_R internalization and D_1_R-H_3_R disruption. In addition, pre-treatment with H_3_R ligands can block D_1_R-induced cell death and prevent D_1_R-H_3_R loss. Finally, the effect of TAT-peptide analogues of D_1_R transmembrane domains in D_1_R-H_3_R stability and function demonstrate that we are observing specific D_1_R-H_3_R, and not D_2_R-H_3_R, signaling and function. Thioperamide has recently been suggested to act via the H_4_R receptor, however, we received similar effects using the H_3_R antagonist VUF 5681 and lost any effects in cells where H_3_R expression was silenced, arguing that the effects are due to D_1_R-H_3_R heteromers.

Besides striatal and cortical cell death, growing evidence points to neuronal dysfunction as responsible for the earliest HD disturbances in cognitive and behavioral changes (6;71). Despite these early changes, no effective treatments are currently available to treat cognitive decline in HD. Moreover, the timing of intervention is also critical, since atrophy and dysfunction progress with age and treatment may be different according to the stage of illness. In this scenario, and given the well-known role of both dopamine and histamine in synaptic plasticity and memory (72–80), it is possible that the therapeutic potential of H_3_R ligands as modulators of D_1_R-H_3_R heteromers could also be extended to improve learning impairments and cognitive decline in HD. This is supported by our data showing that chronic treatment with the H_3_R antagonist thioperamide at 5 months of age prevented motor learning deficits, as well as impaired spatial and recognition memories in mutant Hdh^Q7/Q111^ mice. Importantly, thioperamide treatment does not induce off-target effects (such as alterations in spontaneous locomotor activity or anxiety-like behaviors) neither in wild-type Hdh^Q7/Q7^ nor in mutant Hdh^Q7/Q111^ mice. In addition, early chronic treatment with thioperamide prevented disruption of the heteromer at 6 and 8 months of age and the subsequent cognitive decline. It seems unlikely that there is a direct link between D_1_R-H_3_R heteromers and cognitive deficits, but the data do suggest that whatever neuronal changes occur during progression of the disease they are blocked or at minimum delayed. Importantly, we can say that D_1_R-H_3_R heteromers are required for this effect as thioperamide treatment at 7 months of age (when the heteromer is lost in HD mice) is not able to prevent cognitive and motor learning deficits. This latter result might explain the results of the effects that GSK189254, an H_3_R antagonist, have in a Q175 mouse model of HD (81). The authors saw no change in motor performance and mild improvement in exploratory behavior as measured in the Open Field test and in cognitive function as measured by a T-maze. Our data suggest that D_1_R-H_3_R heteromer expression is crucial to the efficacy of H_3_R antagonists as a therapeutic option in HD.

What disease-driven neuronal changes are prevented by H_3_R antagonism through the D_1_R-H_3_R heteromer is not completely clear. However, we did find that chronic thioperamide treatment at early stages completely rescue the reduction in the density of spinophilin-immunoreactive puncta in HD mice in both hippocampal and cortical areas, suggesting that adequate dopaminergic signaling is required for normal forms of synaptic structural plasticity and cognitive processes. Substantial data support the importance of dopamine receptors for synaptic plasticity in the cortex and hippocampus (82–84). In this view, any dopamine imbalance with both suboptimal and supra-optimal dopamine activity has been reported to modify cognitive performance (85;86). As the early stages of HD may reflect a hyperdopaminergic stage (7;87), treatments reducing dopamine signaling may have therapeutic benefits. In fact, dopamine-depleting drugs such as tetrabenazine or dopamine-stabilizers as pridopidine showed neuroprotective effects in HD mice (88), and improve motor coordination abnormalities in HD patients (12;89), while specific D_1_R inhibition rescues electrophysiological changes in excitatory and inhibitory synaptic transmission in full-length HD mouse models (18). However, none of these treatments have demonstrated cognitive improvements. The suggestion that D_1_R-H_3_R heteromers may be legitimate targets for the treatment of HD shines a spotlight on what continues to be an elusive drug target. Indeed, in the context of this study, the loss of the heteromer in disease progression despite the fact that the receptors themselves are still expressed and functional, points to the heteromers as optimal targets rather than the single receptors. The concept of heteromers have been known for over a decade but physiologic examples have only recently come to be appreciated (33;37;90-95). In sum, our study showing that H_3_R antagonists can prevent learning and memory deficits by blocking D_1_R in D_1_R-H_3_R complexes, along with the role of these heteromers on neuronal cell death, predict a critical role of the histaminergic system as modulator of the dopamine imbalance in HD, and may help to overcome the deleterious effects of directly manipulating DA-production and/or signaling, thus opening new and important alternatives for HD therapeutics.

## Material and Methods

### Human brain slices

Paraffin-embedded *post-mortem* 4 μm-thick brain sections containing caudate-putamen were obtained and provided by the Tissue Bank at Hospital Universitario Fundación Alcorcón (Madrid, Spain) and the Netherlands Brain Bank (Amsterdam, The Netherlands) according to the standardized procedures of both institutions. The samples analyzed were from patients with HD (1 grade 0; 1 grade 1; 2 grade 2; 3 grade 3 and 3 grade 4 patients) or from age matched controls with no neurological disease (3 subjects). All protocols were approved by the institutional ethic committees.

### Cell cultures

Conditionally immortalized wild-type STHdh^Q7^ and mutant STHdh^Q111^ striatal neuronal progenitor cell lines expressing endogenous levels of normal and mutant huntingtin with 7 and 111 glutamines, respectively, have been described previously (96). These cells do not exhibit amino-terminal inclusions allowing the study of changes involved in early HD pathogenesis (96). Striatal cells were grown at 33°C in DMEM (Sigma-Aldrich), supplemented with 10% fetal bovine serum (FBS), 1% streptomycinpenicillin, 2 mM L-glutamine, 1 mM sodium pyruvate, and 400 g/ml G418 (Geneticin; Invitrogen).

HEK-293T cells were grown in Dulbecco’s modified Eagle’s medium (DMEM) (Gibco, Paisley, Scotland, UK) supplemented with 2 mM L-glutamine, 100 μg/ml sodium pyruvate, 100 U/ml penicillin/streptomycin, essential medium non-essential amino acids solution (1/100) and 5% (v/v) heat inactivated fetal bovine serum (Invitrogen, Paisley, Scotland, UK) and were maintained at 37°C in an atmosphere with 5% CO2. Cells were transiently transfected with the corresponding fusion protein cDNA using Lipofectamine 3000 (Invitrogen, Paisley, Scotland, UK).

### Animal models of HD

Knock-in mice, with targeted insertion of 109 CAG repeats that extends the glutamine segment in murine huntingtin to 111 residues, and the corresponding littermates having 7 glutamine residues were maintained on a C57BL/6 genetic background (97). Hdh^Q7/Q111^ heterozygous males and females were intercrossed to generate age-matched Hdh^Q7/Q111^ heterozygous and Hdh^Q7/Q7^ wild-type littermates. Only males were used for all experiments. Hemizigous male mice transgenic for exon 1 of the human huntingtin gene with a greatly expanded CAG repeat (∼115 CAG repeats in R6/1 mice and ∼160 CAG repeats in R6/2 mice) (98) and wild-type littermates were used when indicated in proximity ligation assays. Animals were housed under a 12 h light/dark cycle with food and water ad libitum. All procedures were carried out in accordance with the National Institutes of Health and were approved by the local animal care committee of the Universitat de Barcelona (99/01) and the Generalitat de Catalunya (00/1094) or the Universidad Complutense de Madrid in accordance with the directives of the European Commission.

### Mouse brain slices preparation

For PLA experiments, 2-, 4-, 6- and 8-month-old Hdh^Q7/Q7^ and Hdh^Q7/Q111^ mice were deeply anesthetized and immediately perfused transcardially with saline (PBS) followed by 4% paraformaldehyde (PFA)/phosphate buffer. Brains were removed and post-fixed overnight in the same solution, cryoprotected by immersion in 10, 20, 30% gradient sucrose (24 hours for each sucrose gradient) at 4°C and then frozen in dry ice-cooled methylbutane. Serial coronal cryostat sections (30µm) through the whole brain were collected in PBS-0.025% azide as free-floating sections and stored at 4°C until PLA experiments were performed. For cell death determination, Hdh^Q7/Q111^ and Hdh^Q7/Q7^ mice were killed by cervical dislocation at the age of 4, 5 and 8 months. Mouse brains were rapidly removed and placed in ice-cold oxygenated (O_2_/CO_2_: 95%/5%) Krebs-HCO_3_^-^ buffer (124 mM NaCl, 4 mM KCl, 1.25 mM NaH_2_PO_4_, 1.5 mM MgSO_4_, 1.5 mM CaCl_2_, 10 mM glucose and 26 mM NaHCO_3_, pH 7.4). Cerebral hemisferes were split and sliced coronally using a McIlwain chopper (Ted Pella, Inc, California) in sterile conditions. Striatum, cortex and hippocampal slices (300 µm thick) were kept at 4°C in Krebs-HCO_3_ buffer during the dissection and transferred into a Millicell Insert (Millipore).

### Cell death determination in striatal cells and in mouse organotypic slice cultures

Striatal STHdh^Q7^ or STHdh^Q111^ cells were grown to reach 50 % of confluence on 12-well plates containing 3 cm^2^-glass coverslips. Medium was then replaced by a new supplemented medium containing 0.5 % FBS. Vehicle, SCH 23390, thioperamide or SB 203580 were added at the indicated concentrations to cells and incubated for 1 h before the addition of D_1_R. When TAT-TM peptides were applied to cell cultures, these were added 4 h before the addition of D_1_R agonist. After agonist addition, an additional incubation period of 24 h was performed. Then cells were washed twice in cold-PBS and fixed with 4 % paraformaldehyde for 1 h at 4°C. Sample nuclei were stained with Hoechst 1:1000. Stained cells were then washed with PBS and mounted under glass coverslips with Mowiol. A minimum of 10 fields were taken from each coverslip using a fluorescence microscope and the plugin Image-based Tool for Counting Nuclei for ImageJ was used for the quantification of the total nuclei. In mouse organotypic cultures, brain slices (300 μm thickness, see above) were cultured for 24 h into a Millicell Insert in Neurobasal medium supplemented with 20 % horse serum, 0.5% B27, 2 mM L-glutamine, 100 µg/ml sodium pyruvate, non-essential amino acids solution (1/100) and 100 units/ml penicillin/streptomycin (all supplements were from Invitrogen, Paisley, Scotland, UK) before replacing with fresh medium. Vehicle, SCH 23390, thioperamide were added at the indicated concentrations to organotypic cultures and incubated for 1 h before the addition of D_1_R agonist. TAT-TM peptides were applied to cell cultures 4 h before the addition of D_1_R agonist. After agonist addition, an additional incubation period of 48 h was performed. Then, 10μM propidium iodide (PI) was added to organotypic cultures and maintained at 37°C for 1 h. Organotypic cultures were washed twice in cold-PBS and fixed with 4 % paraformaldehyde for 1 h at 4°C. Total nuclei were stained with Hoechst 1:1000. The Hoechst stained and PI positive nuclei in organotypic cultures were counted to evaluate cell death in the brain slices. Quantification was performed using Leica SP2 confocal microscope (20x; UV, 561 lasers) and the quantification performed with the program Image-based Tool for Counting Nuclei for ImageJ. Cell death is expressed as the percentage of PI positive cells in the total Hoechst-stained nuclei.

### Lentivirus production and cell transduction

Silencing lentiviral vectors were produced by co-transfecting HEK293T producing cells with lentiviral silencing plasmids GIPZ Human histamine H3 receptor shRNA (Clone V3LHS_638095 or Clone V3LHS_638091, Thermo Scientific) with packing plasmid psPAX2 and envelope coding plasmid pMD2.G (Addgene#12260 and #12259, respectively) using the calcium phosphate method. For production of control non silencing lentiviral particles the H_3_R silencing plasmid were substituted with GIPZ Non-silencing Lentiviral shRNA Control (RHS4346, Thermoscientific). Infectious lentiviral particles were harvested at 48 h post-transfection, centrifuged 10 minutes at 900 g to get rid of cell debris, and then filtered through 0.45 μm cellulose acetate filters. The titer of recombinant lentivirus was determined by serial dilution on HEK293T cells. For lentivirus transduction, striatal cells were subcultured to 50% confluence, cells were transduced with H_3_R-shRNA-expressing lentivirus obtained with plasmid (Clone V3LHS_638095) or control-shRNA-expressing lentivirus (LV control) at a multiplicity of infection (MOI) of 10 in the presence of polybrene 5 µg/ml. Virus-containing supernatant was removed after 3 h. Puromycin was added to the culturing media at the final concentration of 1 µg/ml 2 days after infection. 5 days after puromycin selection cells were transduced with the second H_3_R-shRNA-expressing lentivirus obtained with plasmid Clone V3LHS_638091 to improve the level of silencing achieved. LV control infected cells were re-infected with control-shRNA-expressing lentivirus. The second infection was carried out as the first one. Cells were tested 72 h after the second transduction was performed.

### RNA and real-time PCR

RNA was extracted using TRIzol Reagent (Molecular Research Center). 10 μg of total RNA were treated with RQ1 RNAse free DNAse (Promega) according to manufacturer instruction. DNAse treated DNA was quantified again and cDNA was synthesized using 2 μg total RNA with a High Capacity cDNA Reverse Transcription Kit; (Applied Biosystems). The mRNAs of actin, H3R and D1R were amplified by real-time (RT)-PCR using 1 μL cDNA and power SYBER green PCR Master Mix (Applied Biosystems) on a 7500 Real Time PCR system (Applied Biosystems). Primer sequences are as follows: MsACT For: ATGAGCTGCCTGACGGCCAGGTCAT, MsACT Rev: TGGTACCACCAGACAGCAC TGTGTT, H_3_R For: GCAACGCGCTGGTCATGCTC, H_3_R Rev: CCCCGGCCAAAGGTCCAACG, D_1_R FOR: ACCTCTGTGTGATCAGCGTG, AND D_1_R REV: GCGTATGTCCTGCTCAACCT. Thermal cycling conditions for amplification were set at 50°C for 2 min and 95°C for 10 min, respectively. PCR denaturing was set at 95°C for 15 s and annealing/extending at 60°C for 60 s for 40 cycles. mRNA levels normalized for actin are expressed as fold change relative to control cells. The results were quantified with the comparative *C*_t_ method (known as the 2^−δδCt^ method).

### In Situ Proximity Ligation Assays (PLA)

Cells or mouse or human brain slices were mounted on glass slides and treated or not with the indicated concentrations of receptor ligands or TAT-TM peptides for the indicated time. Then, cells or slices were thawed at 4°C, washed in 50 mM Tris-HCl, 0.9% NaCl pH 7.8 buffer (TBS), permeabilized with TBS containing 0.01% Triton X-100 for 10 min and successively washed with TBS. Heteromers were detected using the Duolink II in situ PLA detection Kit (OLink; Bioscience, Uppsala, Sweden) following the instructions of the supplier. A mixture of equal amounts of the primary antibodies: guinea pig anti-D_1_R antibody (1/200 Frontier Institute, Ishikari, Hokkaido, Japan) and rabbit anti-H_3_R antibody (1:200, Alpha diagnostic, San Antonio, Texas, USA) were used to detect D_1_R-H_3_R heteromers together with PLA probes detecting guinea pig or rabbit antibodies, Duolink II PLA probe anti-guinea pig minus and Duolink II PLA probe anti-rabbit plus. Then samples were processed for ligation and amplification with a Detection Reagent Red and were mounted using a DAPI-containing mounting medium. Samples were observed in a Leica SP2 confocal microscope (Leica Microsystems, Mannheim, Germany) equipped with an apochromatic 63X oil-immersion objective (N.A. 1.4), and a 405 nm and a 561 nm laser lines. For each field of view a stack of two channels (one per staining) and 9 to 15 Z stacks with a step size of 1 μm were acquired. For PLA with brain slices, after image processing, the red channel was depicted in green color to facilitate detection on the blue stained nucleus and maintaining the color intensity constant for all images. A quantification of cells containing one or more spots versus total cells (blue nucleus) and, in cells containing spots, the ratio r (number of red spots/cell containing spots) were determined, using the Fiji package (http://pacific.mpi-cbg.de/), considering a total of 600-800 cells from 4-10 different fields within each brain region from 3 different mice per group or from 3 human control subjects, 3 human grade 3 or grade 4 HD patients, 2 grade 0 or grade 1 HD patients or 1 grade 2 HD patient. Nuclei and spots were counted on the maximum projections of each image stack. After getting the projection, each channel was processed individually. The nuclei were segmented by filtering with a median filter, subtracting the background, enhancing the contrast with the Contrast Limited Adaptive Histogram Equalization (CLAHE) plug-in and finally applying a threshold to obtain the binary image and the regions of interest (ROI) around each nucleus. Red spots images were also filtered and thresholded to obtain the binary images. Red spots were counted in each of the ROIs obtained in the nuclei images.

### Membrane preparation and radioligand binding

Striatal cells or mouse striatal, cortical or hippocampal tissue were homogenized in 50 mM Tris-HCl buffer, pH 7.4, containing a protease inhibitor mixture (1/1000, Sigma). The cellular debris was removed by centrifugation at 13, 000 g for 5 min at 4°C, and membranes were obtained by centrifugation at 105, 000 g for 1 h at 4 °C. Membranes were washed three more times at the same conditions before use. Ligand binding was performed with membrane suspension (0.2 mg of protein/ml) in 50 mM Tris–HCl buffer, pH 7.4 containing 10 mM MgCl_2_, at 25°C. To obtain saturation curves, membranes were incubated with increasing free concentrations of [^3^H] SCH 23390 (0.02 nM to 10 nM, PerkinElmer, Boston, MO, USA) or [^3^H]R-a-methyl histamine (0.1 nM to 20 nM [^3^H]RAMH, Amersham, Buckinghamshire, UK) providing enough time to achieve stable equilibrium for the lower ligand concentrations. Nonspecific binding was determined in the presence of 30 µM non-labeled ligand. Free and membrane bound ligand were separated by rapid filtration of 500 µl aliquots in a cell harvester (Brandel, Gaithersburg, MD, USA) through Whatman GF/C filters embedded in 0.3% polyethylenimine that were subsequently washed for 5 s with 5 ml of ice-cold Tris–HCl buffer. The filters were incubated overnight with 10 ml of Ecoscint H scintillation cocktail (National Diagnostics, Atlanta, GA, USA) at room temperature and radioactivity counts were determined using a Tri-Carb 1600 scintillation counter (PerkinElmer, Boston, MO, USA) with an efficiency of 62%. Protein was quantified by the bicinchoninic acid method (Pierce Chemical Co., Rockford, IL, USA) using bovine serum albumin dilutions as standard. Monophasic saturation curves were analyzed by non-linear regression, using the commercial Grafit software (Erithacus Software), by fitting the binding data to the equation previously deduced (equation (3) in (99).

### Immunocytochemistry

Cells (60% confluence) were treated with vehicle or 30 µM SKF 81297 and after 45 min cells were kept at 4 °C to block endocytosis/exocytosis, washed twice in cold-PBS, fixed in 4% paraformaldehyde for 15 min and washed with PBS containing 20 mM glycine (buffer A) to quench the aldehyde groups. After permeabilization with buffer A containing 0.05% Triton X-100 for 5 min, cells were washed with buffer A containing 1% bovine serum albumin (blocking solution) for 1 h and labeled with the primary guinea pig anti-D_1_R antibody (1/100, Frontier Institute, Ishikari, Hokkaido, Japan, ON at 4°C), washed with blocking solution, and stained with the secondary goat Alexa Fluor 488 anti-guinea pig antibody (1:100, Jackson Immunoresearch Laboratories, West Grove, PA, USA, 2 h at RT). Samples were washed twice with blocking solution, once with buffer A and finally with PBS. Nuclei were stained with 1:1000 Hoechst. Cells were mounted with Mowiol and observed in a Leica SP2 confocal microscope.

### Signaling in striatal cells

To determine ERK 1/2 phosphorylation, cells (35, 000/well) were cultured with a non-supplemented medium overnight before pre-treated at 25°C for 20 min with the antagonists and stimulated for an additional 7 min with the indicated agonists. Phosphorylation was determined by alpha-screen bead-based technology using the Amplified Luminiscent Proximity Homogeneous Assay kit (PerkinElmer, Waltham, MA, USA) and the Enspire Multimode Plate Reader (PerkinElmer) following the instructions of the supplier. To determine calcium release, striatal cells were transfected with 4 μg of GCaMP6 calcium sensor (100) using lipofectamine 3000. After 48 h, cells were incubated (0.2 mg of protein/ml in 96-well black, clear bottom microtiter plates) with Mg^+2^-free Locke’s buffer pH 7.4 (154 mM NaCl, 5.6 mM KCl, 3.6 mM NaHCO_3_, 2.3 mM CaCl_2_, 5.6 mM glucose and 5 mM HEPES) supplemented with 10 μM glycine. When TAT-TM peptides treatment was performed they were added 1 hour before the addition of receptor ligands at the indicated concentration. Fluorescence emission intensity of GCaMP6s was recorded at 515 nm upon excitation at 488 nm on an EnSpire® Multimode Plate Reader (PerkinElmer, Boston, MO, USA) for 330 s every 5 s and 100 flashes per well. The fluorescence gain was defined as a delta function of ΔF/F(t) = (F(t) – F0)/F0, where F0 is the average fluorescence intensity in the first six measures from the start of recording and F(t) is the fluorescence intensity at a given time and was expressed in %. To determine p38 phosphorylation, striatal cells (80 % confluence) were cultured with a non-supplemented medium 4 h before the addition of the indicated ligand concentration for the indicated time and were lysed with 50 mM Tris-HCl pH 7.4, 50 mM NaF, 150 mM NaCl, 45 mM β-glycerophosphate, 1% Triton X-100, 20 µM phenyl-arsine oxide, 0.4 mM NaVO_4_ and protease inhibitor cocktail. Lysates (20 µg protein) were processed for Western blot a mixture of a rabbit anti-phospho-p38 MAPK (Thr180/Tyr182) antibody (1:1000, Cell Signaling) and a mouse anti-β-tubulin antibody (1:10, 000, Sigma). Bands were visualized by the addition of a mixture of IRDye 680 anti-rabbit antibody (1:10, 000, Sigma) and IRDye 800 anti-mouse antibody (1:10, 000, Sigma) for 2 h at room temperature and scanned by the Odyssey infrared scanner (LI-COR Biosciences). Band densities were quantified using the Odyssey scanner software. The level of phosphorylated p38 MAPK was normalized for differences in loading using the β-tubulin band intensities.

### Mice thioperamide treatment

Thioperamide maleate salt (Sigma-Aldrich, St. Louis, USA) was prepared fresh daily being dissolved in sterile 0, 9% saline (NaCl) in order to deliver a final dose of 10 mg/kg in a final volume of 0.01 ml/g of body weight, as previously described (101). The vehicle treatment consisted of an equal volume of saline solution. All injections were given via the intra-peritoneal route (*i.p*). Three *i.p* injections per week were administered to wild-type Hdh^Q7/Q7^ and mutant knock-in Hdh^Q7/Q111^ mice from 5 months of age until 6 months of age (when one cohort of animals was perfused to analyze PLA after behavioral assessment) or until 8 months of age (when a second cohort of animals were perfused to analyze PLA at this more advanced disease stage). A total of 11 saline-Hdh^Q7/Q7^ mice, 10 thioperamide-Hdh^Q7/Q7^ mice, 7 saline-Hdh^Q7/Q111^ mice and 9 thioperamide-Hdh^Q7/Q111^ mice were treated. For these experiments, a total of 11 saline-Hdh^Q7/Q7^ mice, 10 thioperamide-Hdh^Q7/Q7^ mice, 7 saline-Hdh^Q7/Q111^ mice and 9 thioperamide-Hdh^Q7/Q111^ mice were treated. Similarly, three *i.p* injections per week were administered to wild-type Hdh^Q7/Q7^ and mutant knock-in Hdh^Q7/Q111^ mice from 7 months of age until 8 months of age to perform the behavioral studies when the D_1_R-H_3_R heteromers were lost. For these experiments, a total of 11 saline-Hdh^Q7/Q7^ mice, 12 thioperamide-Hdh^Q7/Q7^ mice, 10 saline-Hdh^Q7/Q111^ mice and 11 thioperamide-Hdh^Q7/Q111^ mice were treated. All treatments were performed in the afternoon to avoid the stress caused by the treatments during the behavioral assessment. Thus, during behavioral analysis treatments were performed after the evaluation of motor learning or cognitive tasks.

## Behavior assays

Accelerating rotarod was performed as previously described (44). Animals were placed on a motorized rod (30mm diameter). The rotation speed gradually increased from 4 to 40 rpm over the course of 5 min. The time latency was recorded when the animal was unable to keep up on the rotarod with the increasing speed and fell. Rotarod training/testing was performed as 4 trials per day during 3 consecutive days. A resting period of one hour was left between trials. The rotarod apparatus was rigorously cleaned with ethanol between animal trials in order to avoid odors.

For T-maze spontaneous alternation task (T-SAT), the T-maze apparatus used was a wooden maze consisting of three arms, two of them situated at 180° from each other, and the third, representing the stem arm of the T, situated at 90° with respect to the other two. All arms were 45 cm long, 8 cm wide and enclosed by a 20 cm wall. Two identical guillotine doors were placed in the entry of the arms situated at 180°. In the training trial, one arm was closed (new arm) and mice were placed in the stem arm of the T (home arm) and allowed to explore this arm and the other available arm (old arm) for 10 min, after which they were returned to the home cage. After 5 h (LTM), mice were placed in the stem arm of the T-maze and allowed to freely explore all three arms for 5 min. The arm preference was determined by calculating the time spent in each arm x 100/time spent in both arms (old and new arm). The T-maze was rigorously cleaned with ethanol between animal trials in order to avoid odors.

Novel object recognition test (NORT) consisted in a white circular arena with 40 cm diameter and 40 cm high. Mice were first habituated to the open field arena in the absence of objects (2 days, 15 min/day). During these two days of habitation, several parameters were measured to ensure the proper habituation of all mice in the new ambient. As a measure of anxiety or motivation behaviors, the distance that each mice rove in the periphery or in the center of the open field arena was measured as the rove distance in the periphery or in the center x 100/the total distance. The same analysis was performed by counting the number of entries in the periphery and in the center as well as the time that each mouse spent exploring the periphery or the center. The total distance that each mice rove during these two days of habituation was also recorded as a measure to evaluate spontaneous locomotor activity. On the third day, two similar objects were presented to each mouse during 10 min (A, A’ condition) after which the mice were returned to their home cage. Twenty-four hours later (LTM), the same animals were re-tested for 5 min in the arena with a familiar and a new object (A, B condition). The object preference was measured as the time exploring each object × 100/time exploring both objects. The arena was rigorously cleaned with ethanol between animal trials in order to avoid odors. Animals were tracked and recorded with SMART junior software (Panlab, Spain).

### Immunohistochemistry, confocal microscopy and immunofluorescence-positive puncta counting

Saline and thioperamide-treated heterozygous mutant Hdh^Q7/Q111^ and WT Hdh^Q7/Q7^ mice at 6 months of age (n = 3 per group) were deeply anesthetized and immediately perfused transcardially with saline followed by 4% paraformaldehyde (PFA)/ phosphate buffer. Brains were removed and postfixed overnight in the same solution, cryoprotected by immersion in 30% sucrose and then frozen in dry ice-cooled methylbutane. Serial coronal cryostat sections (30 μm) through the whole brain were collected in PBS as free-floating sections. Sections were rinsed three times in PBS and permeabilized and blocked in PBS containing 0.3% Triton X-100 and 3% normal goat serum (Pierce Biotechnology, Rockford, IL) for 15 min at room temperature. The sections were then washed in PBS and incubated overnight at 4°C with Spinophilin (1:250, Millipore) antibody that were detected with Cy3 anti-rabbit secondary antibodies (1:200, Jackson ImmunoResearch, West Grove, PA). As negative controls, some sections were processed as described in the absence of primary antibody and no signal was detected. Confocal microscopy analysis and immunofluorescence-positive puncta counting spinophilin-positive spine-like structures was examined as previously described (44). Briefly, the images were acquired with Zeiss LSM510 META confocal microscope with HeNe lasers. Images were taken using a ×63 numerical aperture objective with ×4 digital zoom and standard (one Airy disc) pinhole. Three coronal sections (30 μm thick) per animal (n=3 per group) spaced 0.24 mm apart containing the motor area M1 or CA1 hippocampus were used. For each slice, we obtained three fields/cortical layer (I, II/III and V) of the M1 area and three fields/CA1 hippocampus (*stratum oriens* and *stratum radiatum*). The number and area of spinophilin-positive puncta were measured using NIH ImageJ version 1.33 by Wayne Rasband (National Institutes of Health, ethesda, MD). To analyze spinophilin immunolabeling, brightness and contrast of fluorescence images were adjusted so that only punctate fluorescence but no weak diffuse background labeling was visible. In the article, we use the term ‘puncta’ and ‘cluster’ interchangeable to refer to discrete points of protein at the fluorescence microscope. Positive puncta/cluster within a specific field was recognized by identifying the presence of overlapping 10–100 pixels.

### Western blot analysis

Saline and thioperamide-treated heterozygous mutant Hdh^Q7/Q111^ and WT Hdh^Q7/Q7^, mice were killed by cervical dislocation at 6 months of age, after behavioral assessment. Brains were quickly removed, dissected, frozen in dry ice and stored at −80°C until use. Protein extraction (n = 5-9 per group, only males) and western blot analysis were performed as previously described (44). The primary antibody 1C2 (1:1000, Millipore) was used. Loading control was performed by reproving the membranes with an antibody to α-actin (1:20.000, MP Biochemicals). ImageJ software was used to quantify the different immunoreactive bands relative to the intensity of the α-actin band in the same membranes within a linear range of detection for the enhanced chemiluminescent kit reagent. Data are expressed as the mean ± SEM of band density.

### Human brain slices

Paraffin-embedded *post-mortem* 4 μm-thick brain sections containing caudate-putamen were obtained and provided by the Tissue Bank at Hospital Universitario Fundación Alcorcón (Madrid, Spain) and the Netherlands Brain Bank (Amsterdam, The Netherlands) according to the standardized procedures of both institutions. The samples analyzed were from patients with HD (1 grade 0; 1 grade 1; 2 grade 2; 3 grade 3 and 3 grade 4 patients) or from age matched controls with no neurological disease (3 subjects). All protocols were approved by the institutional ethic committees.

### Cell cultures

Conditionally immortalized wild-type STHdh^Q7^ and mutant STHdh^Q111^ striatal neuronal progenitor cell lines expressing endogenous levels of normal and mutant huntingtin with 7 and 111 glutamines, respectively, have been described previously (*104*). These cells do not exhibit amino-terminal inclusions allowing the study of changes involved in early HD pathogenesis (*104*). Striatal cells were grown at 33°C in DMEM (Sigma-Aldrich), supplemented with 10% fetal bovine serum (FBS), 1% streptomycinpenicillin, 2 mM L-glutamine, 1 mM sodium pyruvate, and 400 g/ml G418 (Geneticin; Invitrogen).

HEK-293T cells were grown in Dulbecco’s modified Eagle’s medium (DMEM) (Gibco, Paisley, Scotland, UK) supplemented with 2 mM L-glutamine, 100 μg/ml sodium pyruvate, 100 U/ml penicillin/streptomycin, essential medium non-essential amino acids solution (1/100) and 5% (v/v) heat inactivated fetal bovine serum (Invitrogen, Paisley, Scotland, UK) and were maintained at 37°C in an atmosphere with 5% CO2. Cells were transiently transfected with the corresponding fusion protein cDNA using Lipofectamine 3000 (Invitrogen, Paisley, Scotland, UK).

### Animal models of HD

Knock-in mice, with targeted insertion of 109 CAG repeats that extends the glutamine segment in murine huntingtin to 111 residues, and the corresponding littermates having 7 glutamine residues were maintained on a C57BL/6 genetic background (*105*). Hdh^Q7/Q111^ heterozygous males and females were intercrossed to generate age-matched Hdh^Q7/Q111^ heterozygous and Hdh^Q7/Q7^ wild-type littermates. Only males were used for all experiments. Hemizigous male mice transgenic for exon 1 of the human huntingtin gene with a greatly expanded CAG repeat (∼115 CAG repeats in R6/1 mice and ∼160 CAG repeats in R6/2 mice) (*106*) and wild-type littermates were used when indicated in proximity ligation assays. Animals were housed under a 12 h light/dark cycle with food and water ad libitum. All procedures were carried out in accordance with the National Institutes of Health and were approved by the local animal care committee of the Universitat de Barcelona (99/01) and the Generalitat de Catalunya (00/1094) or the Universidad Complutense de Madrid in accordance with the directives of the European Commission.

### Mouse brain slices preparation

For PLA experiments, 2-, 4-, 6- and 8-month-old Hdh^Q7/Q7^ and Hdh^Q7/Q111^ mice were deeply anesthetized and immediately perfused transcardially with saline (PBS) followed by 4% paraformaldehyde (PFA)/phosphate buffer. Brains were removed and post-fixed overnight in the same solution, cryoprotected by immersion in 10, 20, 30% gradient sucrose (24 hours for each sucrose gradient) at 4°C and then frozen in dry ice-cooled methylbutane. Serial coronal cryostat sections (30µm) through the whole brain were collected in PBS-0.025% azide as free-floating sections and stored at 4°C until PLA experiments were performed. For cell death determination, Hdh^Q7/Q111^ and Hdh^Q7/Q7^ mice were killed by cervical dislocation at the age of 4, 5 and 8 months. Mouse brains were rapidly removed and placed in ice-cold oxygenated (O_2_/CO_2_: 95%/5%) Krebs-HCO_3_^-^ buffer (124 mM NaCl, 4 mM KCl, 1.25 mM NaH_2_PO_4_, 1.5 mM MgSO_4_, 1.5 mM CaCl_2_, 10 mM glucose and 26 mM NaHCO_3_, pH 7.4). Cerebral hemisferes were split and sliced coronally using a McIlwain chopper (Ted Pella, Inc, California) in sterile conditions. Striatum, cortex and hippocampal slices (300 µm thick) were kept at 4°C in Krebs-HCO_3_ buffer during the dissection and transferred into a Millicell Insert (Millipore).

### Cell death determination in striatal cells and in mouse organotypic slice cultures

Striatal STHdh^Q7^ or STHdh^Q111^ cells were grown to reach 50 % of confluence on 12-well plates containing 3 cm^2^-glass coverslips. Medium was then replaced by a new supplemented medium containing 0.5 % FBS. Vehicle, SCH 23390, thioperamide or SB 203580 were added at the indicated concentrations to cells and incubated for 1 h before the addition of D_1_R. When TAT-TM peptides were applied to cell cultures, these were added 4 h before the addition of D_1_R agonist. After agonist addition, an additional incubation period of 24 h was performed. Then cells were washed twice in cold-PBS and fixed with 4 % paraformaldehyde for 1 h at 4°C. Sample nuclei were stained with Hoechst 1:1000. Stained cells were then washed with PBS and mounted under glass coverslips with Mowiol. A minimum of 10 fields were taken from each coverslip using a fluorescence microscope and the plugin Image-based Tool for Counting Nuclei for ImageJ was used for the quantification of the total nuclei. In mouse organotypic cultures, brain slices (300 μm thickness, see above) were cultured for 24 h into a Millicell Insert in Neurobasal medium supplemented with 20 % horse serum, 0.5% B27, 2 mM L-glutamine, 100 µg/ml sodium pyruvate, non-essential amino acids solution (1/100) and 100 units/ml penicillin/streptomycin (all supplements were from Invitrogen, Paisley, Scotland, UK) before replacing with fresh medium. Vehicle, SCH 23390, thioperamide were added at the indicated concentrations to organotypic cultures and incubated for 1 h before the addition of D_1_R agonist. TAT-TM peptides were applied to cell cultures 4 h before the addition of D_1_R agonist. After agonist addition, an additional incubation period of 48 h was performed. Then, 10μM propidium iodide (PI) was added to organotypic cultures and maintained at 37°C for 1 h. Organotypic cultures were washed twice in cold-PBS and fixed with 4 % paraformaldehyde for 1 h at 4°C. Total nuclei were stained with Hoechst 1:1000. The Hoechst stained and PI positive nuclei in organotypic cultures were counted to evaluate cell death in the brain slices. Quantification was performed using Leica SP2 confocal microscope (20x; UV, 561 lasers) and the quantification performed with the program Image-based Tool for Counting Nuclei for ImageJ. Cell death is expressed as the percentage of PI positive cells in the total Hoechst-stained nuclei.

### Lentivirus production and cell transduction

Silencing lentiviral vectors were produced by co-transfecting HEK293 producing cellsT with lentiviral silencing plasmids GIPZ Human histamine H3 receptor shRNA (Clone V3LHS_638095 or Clone V3LHS_638091, Thermo Scientific) with packing plasmid psPAX2 and envelope coding plasmid pMD2.G (Addgene#12260 and #12259, respectively) using the calcium phosphate method. For production of control non silencing lentiviral particles the H_3_R silencing plasmid were substituted with GIPZ Non-silencing Lentiviral shRNA Control (RHS4346, Thermoscientific). Infectious lentiviral particles were harvested at 48 h post-transfection, centrifuged 10 minutes at 900 g to get rid of cell debris, and then filtered through 0.45 μm cellulose acetate filters. The titer of recombinant lentivirus was determined by serial dilution on HEK293T cells. For lentivirus transduction, striatal cells were subcultured to 50% confluence, cells were transduced with H_3_R-shRNA-expressing lentivirus obtained with plasmid (Clone V3LHS_638095) or control-shRNA-expressing lentivirus (LV control) at a multiplicity of infection (MOI) of 10 in the presence of polybrene 5 µg/ml. Virus-containing supernatant was removed after 3 h. Puromycin was added to the culturing media at the final concentration of 1 µg/ml 2 days after infection. 5 days after puromycin selection cells were transduced with the second H_3_R-shRNA-expressing lentivirus obtained with plasmid Clone V3LHS_638091 to improve the level of silencing achieved. LV control infected cells were re-infected with control-shRNA-expressing lentivirus. The second infection was carried out as the first one. Cells were tested 72 h after the second transduction was performed.

### RNA and real-time PCR

RNA was extracted using TRIzol Reagent (Molecular Research Center). 10 μg of total RNA were treated with RQ1 RNAse free DNAse (Promega) according to manufacturer instruction. DNAse treated DNA was quantified again and cDNA was synthesized using 2 μg total RNA with a High Capacity cDNA Reverse Transcription Kit; (Applied Biosystems). The mRNAs of actin, H3R and D1R were amplified by real-time (RT)-PCR using 1 μL cDNA and power SYBER green PCR Master Mix (Applied Biosystems) on a 7500 Real Time PCR system (Applied Biosystems). Primer sequences are as follows: MsACT For: ATGAGCTGCCTGACGGCCAGGTCAT, MsACT Rev: TGGTACCACCAGACAGCAC TGTGTT, H_3_R For: GCAACGCGCTGGTCATGCTC, H_3_R Rev: CCCCGGCCAAAGGTCCAACG, D_1_R FOR: ACCTCTGTGTGATCAGCGTG, AND D_1_R REV: GCGTATGTCCTGCTCAACCT. Thermal cycling conditions for amplification were set at 50°C for 2 min and 95°C for 10 min, respectively. PCR denaturing was set at 95°C for 15 s and annealing/extending at 60°C for 60 s for 40 cycles. mRNA levels normalized for actin are expressed as fold change relative to control cells. The results were quantified with the comparative *C*_t_ method (known as the 2^−δδCt^ method).

### In Situ Proximity Ligation Assays (PLA)

Cells or mouse or human brain slices were mounted on glass slides and treated or not with the indicated concentrations of receptor ligands or TAT-TM peptides for the indicated time. Then, cells or slices were thawed at 4°C, washed in 50 mM Tris-HCl, 0.9% NaCl pH 7.8 buffer (TBS), permeabilized with TBS containing 0.01% Triton X-100 for 10 min and successively washed with TBS. Heteromers were detected using the Duolink II in situ PLA detection Kit (OLink; Bioscience, Uppsala, Sweden) following the instructions of the supplier. A mixture of equal amounts of the primary antibodies: guinea pig anti-D_1_R antibody (1/200 Frontier Institute, Ishikari, Hokkaido, Japan) and rabbit anti-H_3_R antibody (1:200, Alpha diagnostic, San Antonio, Texas, USA) were used to detect D_1_R-H_3_R heteromers together with PLA probes detecting guinea pig or rabbit antibodies, Duolink II PLA probe anti-guinea pig minus and Duolink II PLA probe anti-rabbit plus. Then samples were processed for ligation and amplification with a Detection Reagent Red and were mounted using a DAPI-containing mounting medium. Samples were observed in a Leica SP2 confocal microscope (Leica Microsystems, Mannheim, Germany) equipped with an apochromatic 63X oil-immersion objective (N.A. 1.4), and a 405 nm and a 561 nm laser lines. For each field of view a stack of two channels (one per staining) and 9 to 15 Z stacks with a step size of 1 μm were acquired. For PLA with brain slices, after image processing, the red channel was depicted in green color to facilitate detection on the blue stained nucleus and maintaining the color intensity constant for all images. A quantification of cells containing one or more spots versus total cells (blue nucleus) and, in cells containing spots, the ratio r (number of red spots/ cell containing spots) were determined, using the Fiji package (http://pacific.mpi-cbg.de/), considering a total of 600-800 cells from 4-10 different fields within each brain region from 3 different mice per group or from 3 human control subjects, 3 human grade 3 or grade 4 HD patients, 2 grade 0 or grade 1 HD patients or 1 grade 2 HD patient. Nuclei and spots were counted on the maximum projections of each image stack. After getting the projection, each channel was processed individually. The nuclei were segmented by filtering with a median filter, subtracting the background, enhancing the contrast with the Contrast Limited Adaptive Histogram Equalization (CLAHE) plug-in and finally applying a threshold to obtain the binary image and the regions of interest (ROI) around each nucleus. Red spots images were also filtered and thresholded to obtain the binary images. Red spots were counted in each of the ROIs obtained in the nuclei images.

### Membrane preparation and radioligand binding

Striatal cells or mouse striatal, cortical or hippocampal tissue were homogenized in 50 mM Tris-HCl buffer, pH 7.4, containing a protease inhibitor mixture (1/1000, Sigma). The cellular debris was removed by centrifugation at 13,000 g for 5 min at 4°C, and membranes were obtained by centrifugation at 105,000 g for 1 h at 4 °C. Membranes were washed three more times at the same conditions before use. Ligand binding was performed with membrane suspension (0.2 mg of protein/ml) in 50 mM Tris–HCl buffer, pH 7.4 containing 10 mM MgCl_2_, at 25°C. To obtain saturation curves, membranes were incubated with increasing free concentrations of [^3^H] SCH 23390 (0.02 nM to 10 nM, PerkinElmer, Boston, MO, USA) or [^3^H]R-a-methyl histamine (0.1 nM to 20 nM [^3^H]RAMH, Amersham, Buckinghamshire, UK) providing enough time to achieve stable equilibrium for the lower ligand concentrations. Nonspecific binding was determined in the presence of 30 µM non-labeled ligand. Free and membrane bound ligand were separated by rapid filtration of 500 µl aliquots in a cell harvester (Brandel, Gaithersburg, MD, USA) through Whatman GF/C filters embedded in 0.3% polyethylenimine that were subsequently washed for 5 s with 5 ml of ice-cold Tris–HCl buffer. The filters were incubated overnight with 10 ml of Ecoscint H scintillation cocktail (National Diagnostics, Atlanta, GA, USA) at room temperature and radioactivity counts were determined using a Tri-Carb 1600 scintillation counter (PerkinElmer, Boston, MO, USA) with an efficiency of 62%. Protein was quantified by the bicinchoninic acid method (Pierce Chemical Co., Rockford, IL, USA) using bovine serum albumin dilutions as standard. Monophasic saturation curves were analyzed by non-linear regression, using the commercial Grafit software (Erithacus Software), by fitting the binding data to the equation previously deduced (equation (3) in (*107*).

### Immunocytochemistry

Cells (60% confluence) were treated with vehicle or 30 µM SKF 81297 and after 45 min cells were kept at 4 °C to block endocytosis/exocytosis, washed twice in cold-PBS, fixed in 4% paraformaldehyde for 15 min and washed with PBS containing 20 mM glycine (buffer A) to quench the aldehyde groups. After permeabilization with buffer A containing 0.05% Triton X-100 for 5 min, cells were washed with buffer A containing 1% bovine serum albumin (blocking solution) for 1 h and labeled with the primary guinea pig anti-D_1_R antibody (1/100, Frontier Institute, Ishikari, Hokkaido, Japan, ON at 4°C), washed with blocking solution, and stained with the secondary goat Alexa Fluor 488 anti-guinea pig antibody (1:100, Jackson Immunoresearch Laboratories, West Grove, PA, USA, 2 h at RT). Samples were washed twice with blocking solution, once with buffer A and finally with PBS. Nuclei were stained with 1:1000 Hoechst. Cells were mounted with Mowiol and observed in a Leica SP2 confocal microscope.

### Signaling in striatal cells

To determine ERK 1/2 phosphorylation, cells (35, 000/well) were cultured with a non-supplemented medium overnight before pre-treated at 25°C for 20 min with the antagonists, and stimulated for an additional 7 min with the indicated agonists. Phosphorylation was determined by alpha-screen bead-based technology using the Amplified Luminiscent Proximity Homogeneous Assay kit (PerkinElmer, Waltham, MA, USA) and the Enspire Multimode Plate Reader (PerkinElmer) following the instructions of the supplier. To determine calcium release, striatal cells were transfected with 4 μg of GCaMP6 calcium sensor (*108*) using lipofectamine 3000. After 48 h, cells were incubated (0.2 mg of protein/ml in 96-well black, clear bottom microtiter plates) with Mg^+2^-free Locke’s buffer pH 7.4 (154 mM NaCl, 5.6 mM KCl, 3.6 mM NaHCO_3_, 2.3 mM CaCl_2_, 5.6 mM glucose and 5 mM HEPES) supplemented with 10 μM glycine. When TAT-TM peptides treatment was performed they were added 1 hour before the addition of receptor ligands at the indicated concentration. Fluorescence emission intensity of GCaMP6s was recorded at 515 nm upon excitation at 488 nm on an EnSpire® Multimode Plate Reader (PerkinElmer, Boston, MO, USA) for 330 s every 5 s and 100 flashes per well. The fluorescence gain was defined as a delta function of ΔF/F(t) = (F(t) – F0)/F0, where F0 is the average fluorescence intensity in the first six measures from the start of recording and F(t) is the fluorescence intensity at a given time and was expressed in %. To determine p38 phosphorylation, striatal cells (80 % confluence) were cultured with a non-supplemented medium 4 h before the addition of the indicated ligand concentration for the indicated time and were lysed with 50 mM Tris-HCl pH 7.4, 50 mM NaF, 150 mM NaCl, 45 mM β-glycerophosphate, 1% Triton X-100, 20 µM phenyl-arsine oxide, 0.4 mM NaVO_4_ and protease inhibitor cocktail. Lysates (20 µg protein) were processed for Western blot a mixture of a rabbit anti-phospho-p38 MAPK (Thr180/Tyr182) antibody (1:1000, Cell Signaling) and a mouse anti-β-tubulin antibody (1:10, 000, Sigma). Bands were visualized by the addition of a mixture of IRDye 680 anti-rabbit antibody (1:10, 000, Sigma) and IRDye 800 anti-mouse antibody (1:10, 000, Sigma) for 2 h at room temperature and scanned by the Odyssey infrared scanner (LI-COR Biosciences). Band densities were quantified using the Odyssey scanner software. The level of phosphorylated p38 MAPK was normalized for differences in loading using the β-tubulin band intensities.

### Mice thioperamide treatment

Thioperamide maleate salt (Sigma-Aldrich, St. Louis, USA) was prepared fresh daily being dissolved in sterile 0, 9% saline (NaCl) in order to deliver a final dose of 10 mg/kg in a final volume of 0.01 ml/g of body weight, as previously described (*109*). The vehicle treatment consisted of an equal volume of saline solution. All injections were given via the intra-peritoneal route (*i.p*). Three *i.p* injections per week were administered to wild-type Hdh^Q7/Q7^ and mutant knock-in Hdh^Q7/Q111^ mice from 5 months of age until 6 months of age (when one cohort of animals was perfused to analyze PLA after behavioral assessment) or until 8 months of age (when a second cohort of animals were perfused to analyze PLA at this more advanced disease stage). A total of 11 saline-Hdh^Q7/Q7^ mice, 10 thioperamide-Hdh^Q7/Q7^ mice, 7 saline-Hdh^Q7/Q111^ mice and 9 thioperamide-Hdh^Q7/Q111^ mice were treated. For these experiments, a total of 11 saline-Hdh^Q7/Q7^ mice, 10 thioperamide-Hdh^Q7/Q7^ mice, 7 saline-Hdh^Q7/Q111^ mice and 9 thioperamide-Hdh^Q7/Q111^ mice were treated. Similarly, three *i.p* injections per week were administered to wild-type Hdh^Q7/Q7^ and mutant knock-in Hdh^Q7/Q111^ mice from 7 months of age until 8 months of age to perform the behavioral studies when the D_1_R-H_3_R heteromers were lost. For these experiments, a total of 11 saline-Hdh^Q7/Q7^ mice, 12 thioperamide-Hdh^Q7/Q7^ mice, 10 saline-Hdh^Q7/Q111^ mice and 11 thioperamide-Hdh^Q7/Q111^ mice were treated. All treatments were performed in the afternoon to avoid the stress caused by the treatments during the behavioral assessment. Thus, during behavioral analysis treatments were performed after the evaluation of motor learning or cognitive tasks.

## Behavior assays

Accelerating rotarod was performed as previously described (*44*). Animals were placed on a motorized rod (30mm diameter). The rotation speed gradually increased from 4 to 40 rpm over the course of 5 min. The time latency was recorded when the animal was unable to keep up on the rotarod with the increasing speed and fell. Rotarod training/testing was performed as 4 trials per day during 3 consecutive days. A resting period of one hour was left between trials. The rotarod apparatus was rigorously cleaned with ethanol between animal trials in order to avoid odors.

For T-maze spontaneous alternation task (T-SAT), the T-maze apparatus used was a wooden maze consisting of three arms, two of them situated at 180° from each other, and the third, representing the stem arm of the T, situated at 90° with respect to the other two. All arms were 45 cm long, 8 cm wide and enclosed by a 20 cm wall. Two identical guillotine doors were placed in the entry of the arms situated at 180°. In the training trial, one arm was closed (new arm) and mice were placed in the stem arm of the T (home arm) and allowed to explore this arm and the other available arm (old arm) for 10 min, after which they were returned to the home cage. After 5 h (LTM), mice were placed in the stem arm of the T-maze and allowed to freely explore all three arms for 5 min. The arm preference was determined by calculating the time spent in each arm x 100/time spent in both arms (old and new arm). The T-maze was rigorously cleaned with ethanol between animal trials in order to avoid odors.

Novel object recognition test (NORT) consisted in a white circular arena with 40 cm diameter and 40 cm high. Mice were first habituated to the open field arena in the absence of objects (2 days, 15 min/day). During these two days of habitation, several parameters were measured to ensure the proper habituation of all mice in the new ambient. As a measure of anxiety or motivation behaviors, the distance that each mice rove in the periphery or in the center of the open field arena was measured as the rove distance in the periphery or in the center x 100/the total distance. The same analysis was performed by counting the number of entries in the periphery and in the center as well as the time that each mice spent exploring the periphery or the center. The total distance that each mice rove during this two days of habituation was also recorded as a measure to evaluate spontaneous locomotor activity. On the third day, two similar objects were presented to each mouse during 10 min (A, A’ condition) after which the mice were returned to their home cage. Twenty-four hours later (LTM), the same animals were re-tested for 5 min in the arena with a familiar and a new object (A, B condition). The object preference was measured as the time exploring each object × 100/time exploring both objects. The arena was rigorously cleaned with ethanol between animal trials in order to avoid odors. Animals were tracked and recorded with SMART junior software (Panlab, Spain).

### Immunohistochemistry, confocal microscopy and immunofluorescence-positive puncta counting

Saline and thioperamide-treated heterozygous mutant Hdh^Q7/Q111^ and WT Hdh^Q7/Q7^ mice at 6 months of age (n = 3 per group) were deeply anesthetized and immediately perfused transcardially with saline followed by 4% paraformaldehyde (PFA)/ phosphate buffer. Brains were removed and postfixed overnight in the same solution, cryoprotected by immersion in 30% sucrose and then frozen in dry ice-cooled methylbutane. Serial coronal cryostat sections (30 μm) through the whole brain were collected in PBS as free-floating sections. Sections were rinsed three times in PBS and permeabilized and blocked in PBS containing 0.3% Triton X-100 and 3% normal goat serum (Pierce Biotechnology, Rockford, IL) for 15 min at room temperature. The sections were then washed in PBS and incubated overnight at 4°C with Spinophilin (1:250, Millipore) antibody that were detected with Cy3 anti-rabbit secondary antibodies (1:200, Jackson ImmunoResearch, West Grove, PA). As negative controls, some sections were processed as described in the absence of primary antibody and no signal was detected. Confocal microscopy analysis and immunofluorescence-positive puncta counting spinophilin-positive spine-like structures was examined as previously described (*44*). Briefly, the images were acquired with Zeiss LSM510 META confocal microscope with HeNe lasers. Images were taken using a ×63 numerical aperture objective with ×4 digital zoom and standard (one Airy disc) pinhole. Three coronal sections (30 μm thick) per animal (n=3 per group) spaced 0.24 mm apart containing the motor area M1 or CA1 hippocampus were used. For each slice, we obtained three fields/cortical layer (I, II/III and V) of the M1 area and three fields/CA1 hippocampus (*stratum oriens* and *stratum radiatum*). The number and area of spinophilin-positive puncta were measured using NIH ImageJ version 1.33 by Wayne Rasband (National Institutes of Health, ethesda, MD). To analyze spinophilin immunolabeling, brightness and contrast of fluorescence images were adjusted so that only punctate fluorescence but no weak diffuse background labeling was visible. In the article, we use the term ‘puncta’ and ‘cluster’ interchangeable to refer to discrete points of protein at the fluorescence microscope. Positive puncta/cluster within a specific field was recognized by identifying the presence of overlapping 10–100 pixels.

### Western blot analysis

Saline and thioperamide-treated heterozygous mutant Hdh^Q7/Q111^ and WT Hdh^Q7/Q7^, mice were killed by cervical dislocation at 6 months of age, after behavioral assessment. Brains were quickly removed, dissected, frozen in dry ice and stored at −80°C until use. Protein extraction (n = 5-9 per group, only males) and western blot analysis were performed as previously described (*44*). The primary antibody 1C2 (1:1000, Millipore) was used. Loading control was performed by reproving the membranes with an antibody to α-actin (1:20.000, MP Biochemicals). ImageJ software was used to quantify the different immunoreactive bands relative to the intensity of the α-actin band in the same membranes within a linear range of detection for the enhanced chemiluminiscent kit reagent. Data are expressed as the mean ± SEM of band density.

## Author contributions

D.M, M.P, S.G and PJM designed the experiments and wrote the manuscript. D.M performed and analyzed viability, calcium, internalization and organotypic culture experiments. M.P performed all the treatments in mice, conducted and analyzed the behavior tests, obtained all the tissue samples and prepared tissue slices for PLA, organotypic culture and mRNA experiments, performed and analyzed western blot experiments and conducted and analysed spinophilin-immureactive experiments. E.M performed PLA experiments and PLA quantification. M.R assisted with function and viability experiments in cells and organotypic culture. J.B performed the binding experiments and assisted with calcium, internalization and cell death experiments. P.G performed all the shRNA related experiments and conducted and analyzed mRNA experiments. A.Ch helped with the R6 and human PLA experiments, L.A.H designed, synthesized and purified the disrupting peptides, M.S aided with the trafficking experiments, An C and V.C performed and analyzed binding experiments, E.C and S.F aided with the disrupting peptide experiments, M.G provided all the human samples, discussed the results and edited the manuscript. J.A aided with the in vivo experiments and analysis. C.LL designed, supervised experiments, discussed, and helped write the manuscript. S.G and PJM conceived the idea, designed, supervised and coordinated the project, analyzed the results, and wrote the manuscript.

## Acknowledgements

We are very grateful to Ana Lopez for technical assistance. Dr. Teresa Rodrigo and the staff of the animal care facility (Facultat de Psicologia, Universitat de Barcelona), for their support and advice. This work was supported by grants from Ministerio de Economia y Competitividad (SAF2012-39142 to S.G., SAF2011-29507 to J.A., SAF2012-35759 to M.G., and SAF2011-23813 to E.C.,); Centro de Investigacion Biomédicas en Red sobre Enfermedades Neurodegenerativas (CIBERNED CB06/05/0064, CB06/05/0054, CB06/05/0042, and CB06/05/0005); Generalitat de Catalunya, Spain (2014SGR-00968 to J.A. and 2014SGR-1236 to E.C); Grant 20140610 from Fundació La Marató de TV3 to E.C.; RSC Grant Project RG140118, Jerome LeJeune Foundation FJL-01/01/2013, BBSRC BB/N504282/3 and start-up funds from QMUL. We thank Manel Bosch at UB and Paul Thomas at the Henry Welcome Laboratory for Cell Imaging at UEA for their help with the microscopy.

## Supplemental Figure Legends

**Figure S1.**
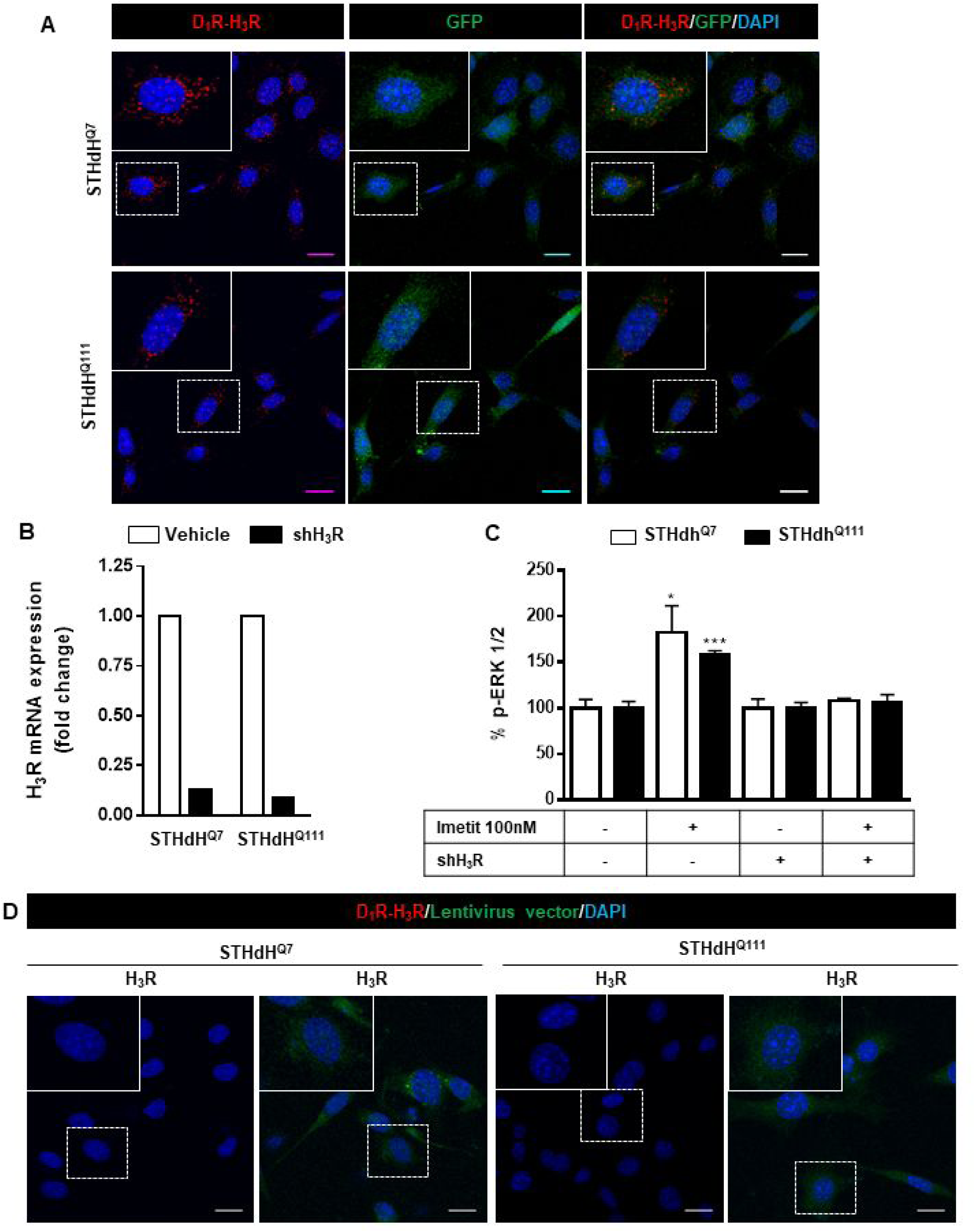
Negative controls for Proximity Ligation Assays (PLA) in striatal cells not depleted or H_3_R depleted by shRNA. In (**A**), Proximity Ligation Assays (PLA) were performed in STHdH^Q7^ and STHdH^Q111^ cells not H_3_R depleted but infected with GIPZ Non-silencing Lentiviral shRNA Control plasmid. D_1_R-H_3_R heteromers were visualized as red spots around blue colored DAPI stained nucleus (left panels), in infected cells stained in green due to the GFP expression included in the plasmid (middle panel). Merge images are given in the right panels. In (**B**), controls showing that H_3_R mRNA is not present in cells depleted of H_3_R by shRNA. STHdH^Q7^ and STHdH^Q111^ cells were not infected or infected with lentiviral silencing plasmid GIPZ Human histamine H3 receptor shRNA (shH_3_R). Values represent fold change respect to non-silencing vector. In (**C**) controls showing the lack of H_3_R stimulated signaling in cells depleted of H_3_R by shRNA. STHdH^Q7^ or STHdH^Q111^ cells were not stimulated (basal) or stimulated with the H_3_R agonist imetit (100 nM) and ERK 1/2 phosphorylation was determined. Values represent mean ± SEM (n = 3) of percentage of phosphorylation relative to basal levels found in untreated cells. Student’s *t* test showed significant differences over basal conditions (*p<0.05, ***p<0.001). In (**D**), PLA were performed in the absence of the D_1_R primary antibody using STHdH^Q7^ or STHdH^Q111^ cells not infected (left panels) or infected (right panels) with GIPZ Non-silencing Lentiviral shRNA Control plasmid. Scale bar: 20 μm.

**Figure S2.**
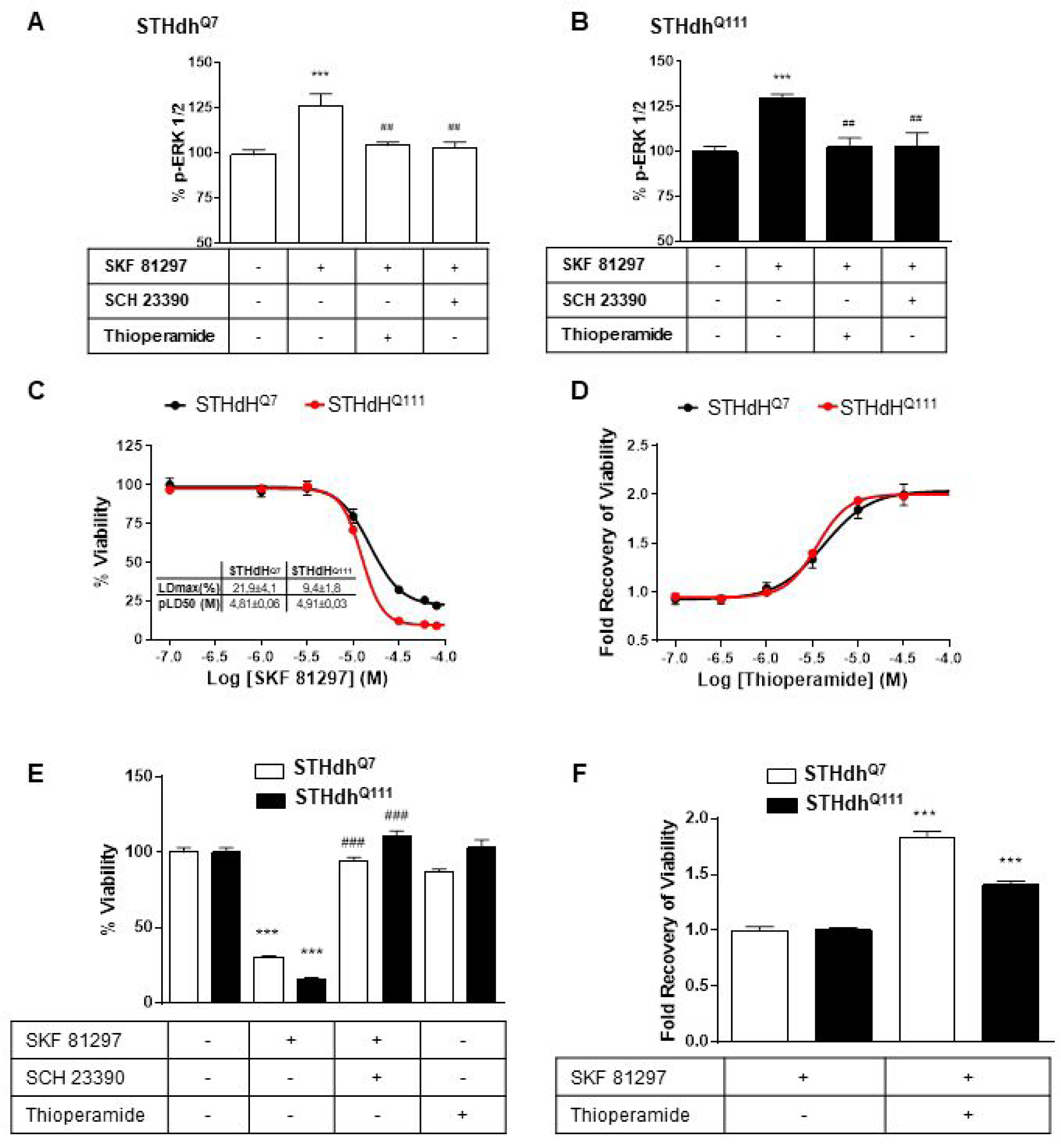
H_3_R ligands revert the D_1_R-mediated decreases in STHdH^Q7^ and STHdH^Q111^ cell viability. STHdH^Q7^ (**A**) or STHdH^Q111^ (**B**) cells were treated for 20 min with vehicle, D_1_R antagonist SCH 23390 (1 μM) or the H_3_R antagonist thioperamide (1 μM) before the addition of SKF 81297 (100 nM) for an additional incubation period of 10 min and ERK 1/2 phosphorylation was determined. Values represent mean ± SEM (n = 3 to 4) of percentage of phosphorylation relative to basal levels found in untreated cells (control). One-way ANOVA followed by Bonferroni post hoc tests showed a significant effect over basal (***p < 0.001) or over SKF 81297 treatment (^##^p < 0.01). In (**C, D**), cell viability was determined in STHdH^Q7^ (black curves) or STHdH^Q111^ cells (red curves) pre-treated for 60 min with vehicle (**C**), or with the H_3_R antagonist thioperamide 10 μM (**B**) prior overstimulation with SKF 81297 (increasing concentrations in **A** or 30 μM in **B**). Values represent mean ± SEM (n = 24 to 30) of percentage of viable cells respect to vehicle-treated cells (**C**) or the cell viability recovery expressed as in-fold respect to SKF 81297 treated cells (**D**). In (**E** and **F**) the effect of D_1_R antagonist, H_3_R antagonist and silencing vector transfection in striatal cells viability is shown. STHdH^Q7^ and STHdH^Q111^ cells were not infected (**E**) or infected (**F**) with GIPZ Non-silencing Lentiviral shRNA Control plasmid. Cells were pretreated for 60 min with vehicle, 10 μM SCH 23390 or 10 μM thioperamide prior over-stimulation with SKF 81297 (30 μM). Values represent mean ± SEM (n = 7 to 22) of percentage of viable cells respect to vehicle-treated cells (**E**) or the cell viability recovery expressed as in-fold respect to SKF 81297 treated cells (**F**). Student’s *t* test showed a significant (***p < 0.001) effect over not treated cells (**E**) or SKF 81297 treated cells (**F**).

**Figure S3.**
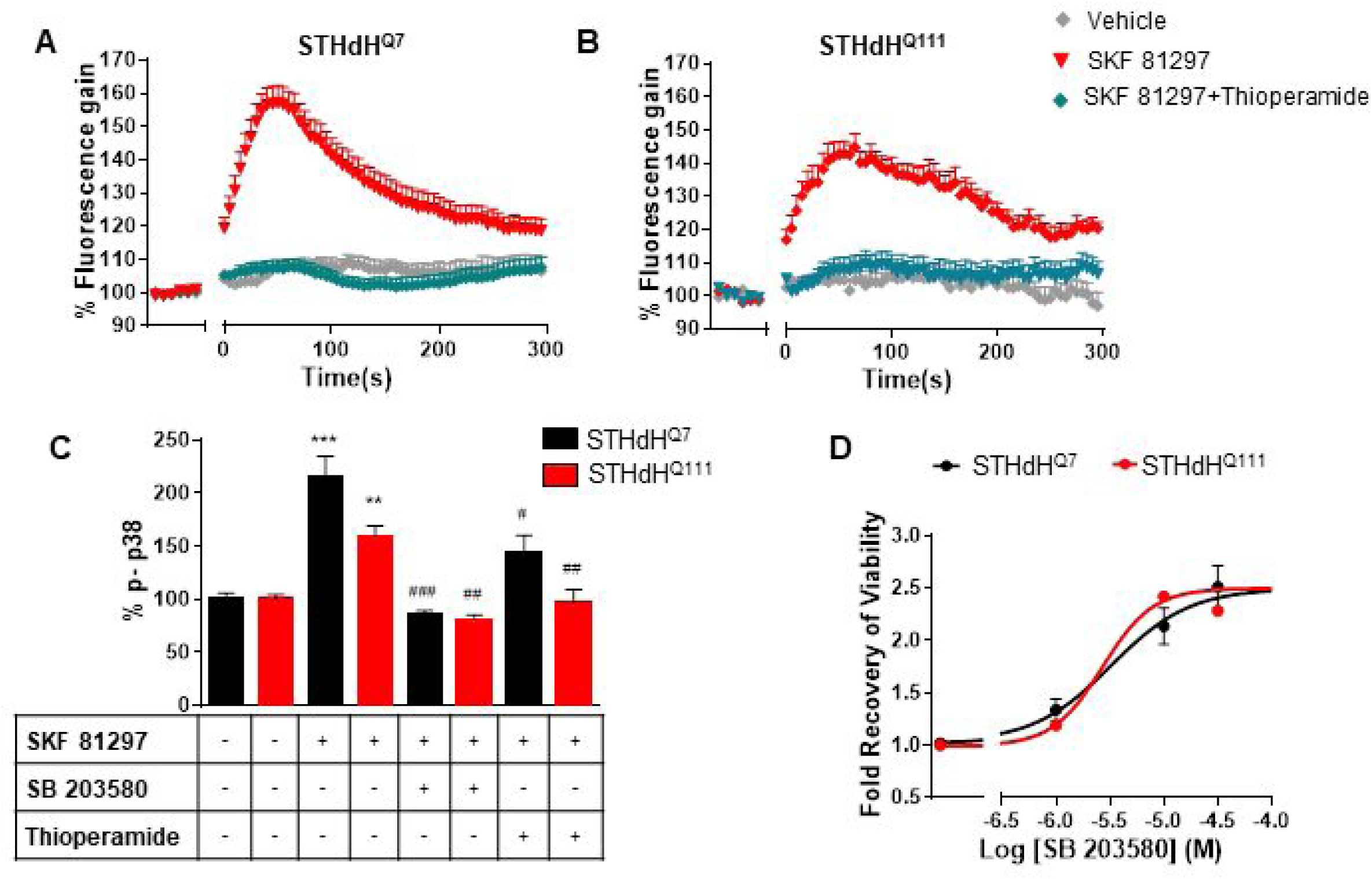
H_3_R ligands revert the D_1_R-mediated decreases in cell viability in STHdH^Q7^ and STHdH^Q111^ by modulating calcium signaling and p38 phosphorylation. In (**A** and **B**), STHdH^Q7^ (**A**) or STHdH^Q111^ (**B**) cells were pre-treated for 20 min with vehicle or with the H_3_R antagonist thioperamide (10 μM) and were not stimulated or overstimulated with SKF 81297 (30 μM) prior intracellular calcium release determination. For each curve values are expressed as a percentage of increase with respect to untreated not overstimulated cells and are mean ± SEM of 3 to 9 independent experiments. In (**C**), STHdH^Q7^ or STHdH^Q111^ cells were treated for 20 min with medium (control), with SB 203580 (10 μM) or with the H_3_R antagonist thioperamide (10 μM). Cells were overstimulated with SKF 81297 (30 μM) and p38 phosphorylation was determined. Values represent mean ± SEM (n = 3) and are expressed as percentage over control. One-way ANOVA followed by Bonferroni post hoc tests showed a significant effect over control (**p < 0.01, ***p < 0.001) or over SKF 81297 treatment (^#^p < 0.05, ^##^p < 0.01, ^###^p < 0.001). In (**D**), cell viability was determined in STHdH^Q7^ (black curves) or STHdH^Q111^ cells (red curves) pre-treated for 60 min with the p38 inhibitor SB 203580 prior overstimulation with SKF 81297 (30 μM). Values represent mean ± SEM (n = 24 to 30) of the cell viability recovery expressed as in-fold respect to SKF 81297 treated cells (**D**).

**Figure S4.**
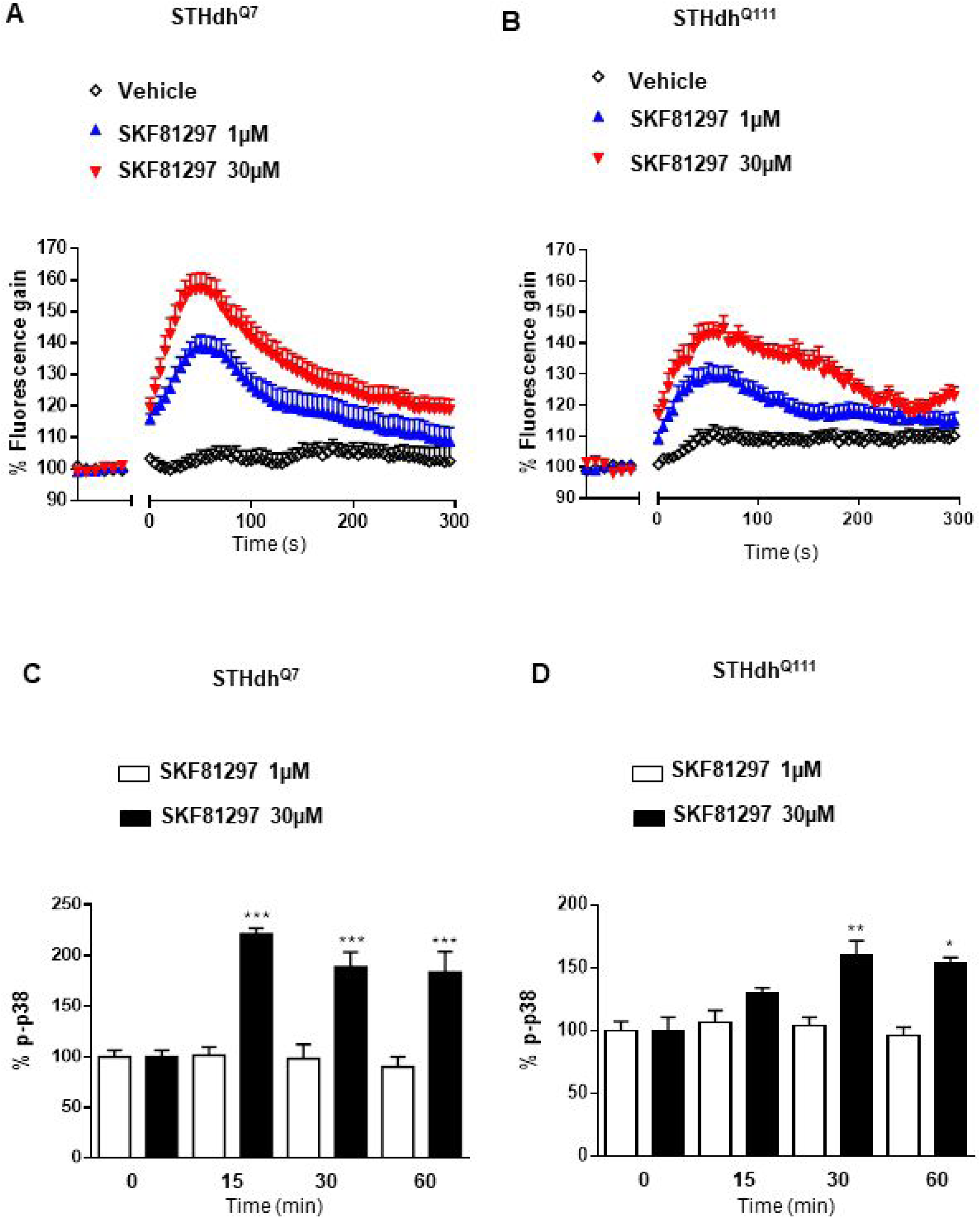
Effect of low and high SKF 81297 concentrations in p-p38 and intracellular calcium release. STHdH^Q7^ (**A** and **C**) and STHdH^Q111^ (**B** and **D**) cells were time-dependent stimulated with 1 μM or 30 μM SKF 81297 and intracellular calcium release (**A** and **B**) or p-p38 phosphorylation (**C** and **D**) was determined. In (**A** and **B**), curves are mean ± SEM of 3 to 6 independent experiments. In (**C** and **D**) values represent mean ± SEM of two independent experiments performed per triplicate of percentage of phosphorylation respect to vehicle-treated cells. Student’s *t* test showed a significant (*p < 0.05, **p < 0.01, ***p < 0.001) effect over not treated cells.

**Figure S5.**
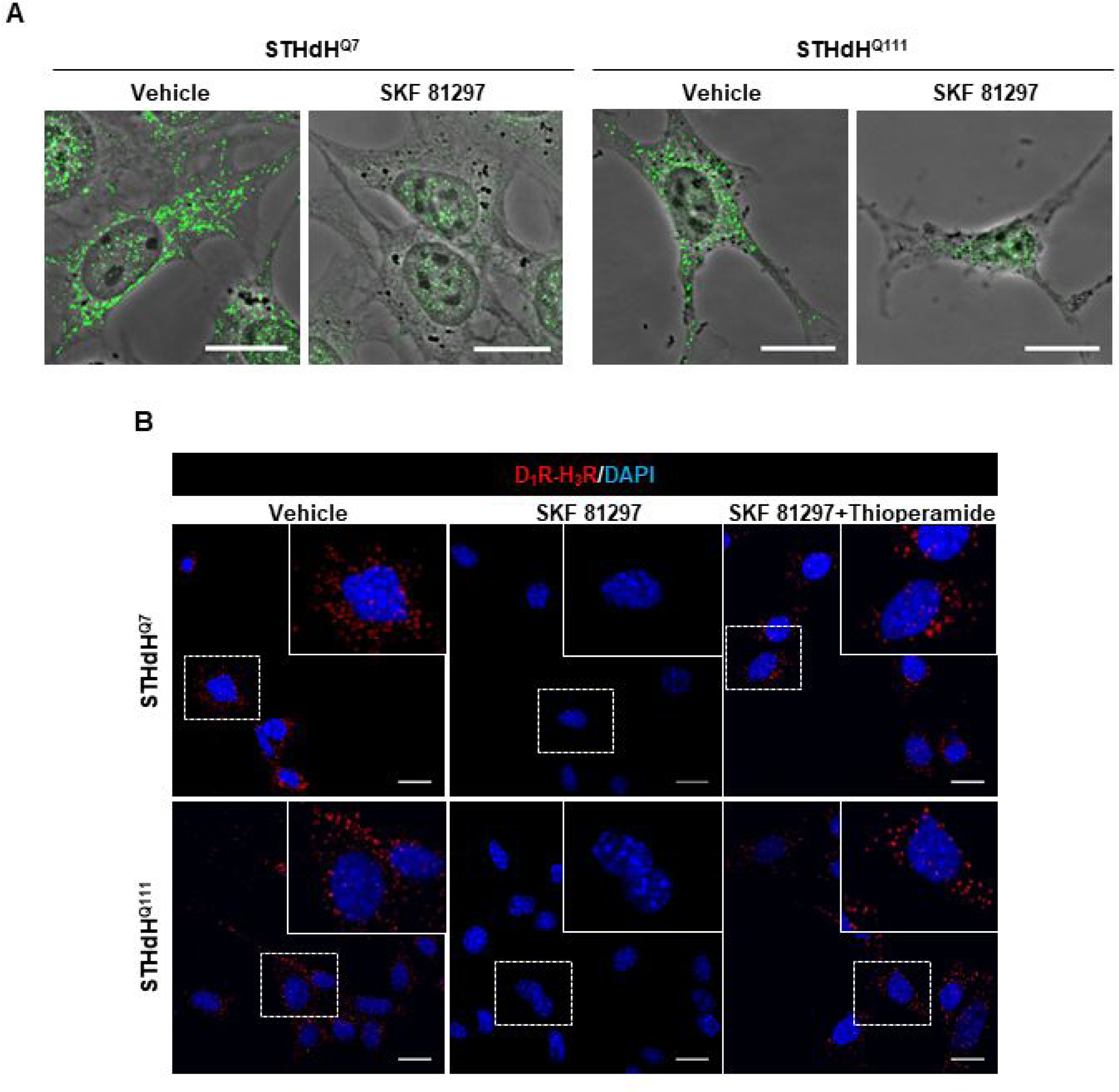
H_3_R ligands revert the D_1_R overstimulation-induced heteromer disruption in striatal cells. In (**A**), superposition of phase contrast and confocal microscopy (superimposed Z stacks) images were shown for immunostained D_1_R (green) in STHdh^Q7^ and STHdh^Q111^ cells treated with vehicle (control) or with SKF 81297 (30 μM) for 45 min. In (**B**), Proximity Ligation Assays (PLA) were performed in STHdH^Q7^ or STHdH^Q111^ cells pre-treated for 60 min with vehicle or with the H_3_R antagonist thioperamide (10µM) before addition of medium (in the case of the vehicle control) or SKF 81297 (30 μM, 45 min). D_1_R-H_3_R heteromers were visualized as red spots around blue colored DAPI stained nucleus in control and H_3_R ligands-treated cells, but not in SKF 81297 only treated cells. Scale bar: 20 μm.

**Figure S6.**
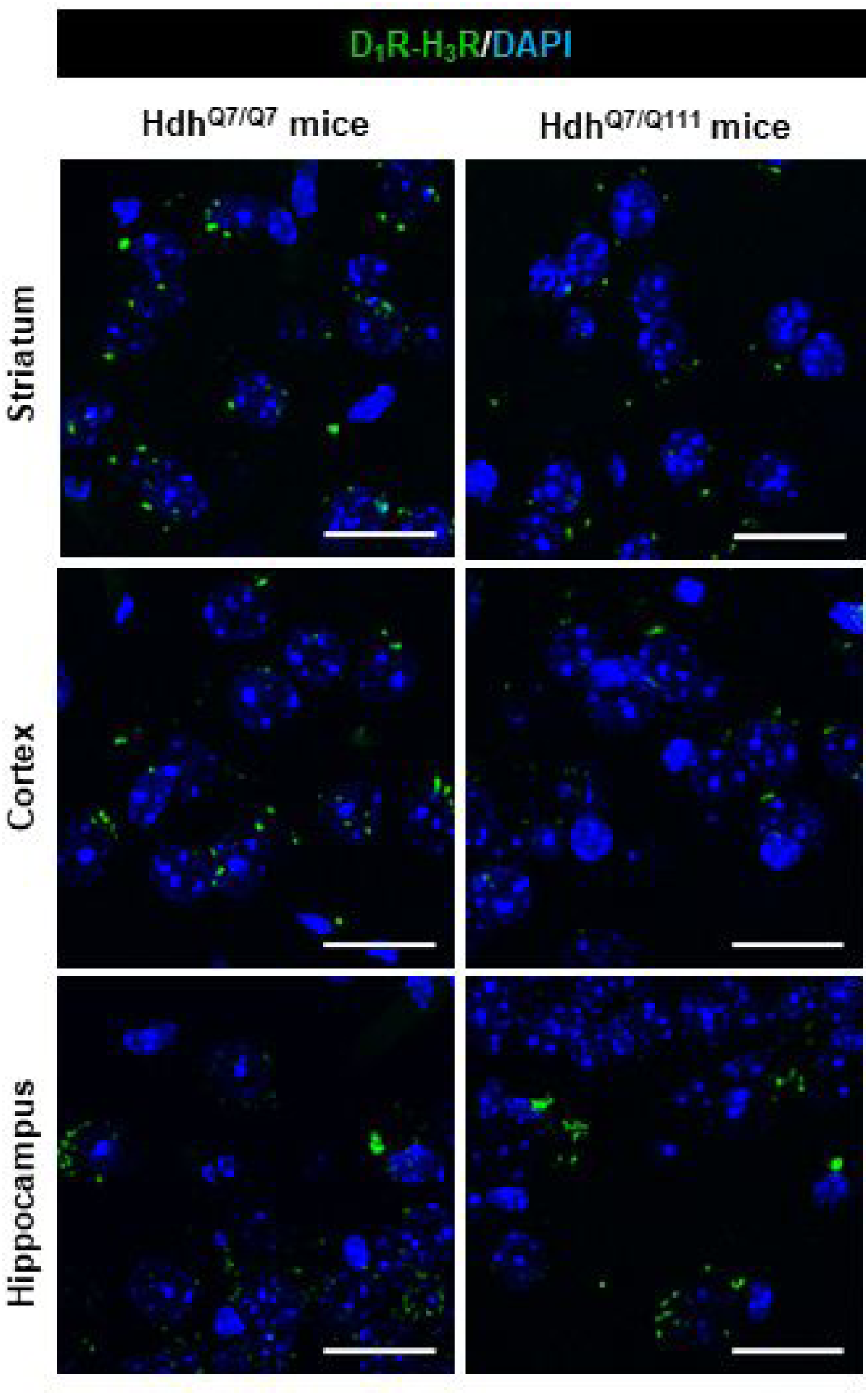
D_1_R-H_3_R heteromer are expressed in 2-month-old Hdh^Q7/Q7^ and Hdh^Q7/Q111^ mice. Proximity Ligation Assays (PLA) were performed using striatal, cortical or hippocampal slices from 2-month-old Hdh^Q7/Q7^ and Hdh^Q7/Q111^ mice. D_1_R-H_3_R heteromers were visualized in all slices as green spots around blue colored DAPI stained nucleus. Scale bar: 20 μm.

**Figure S7.**
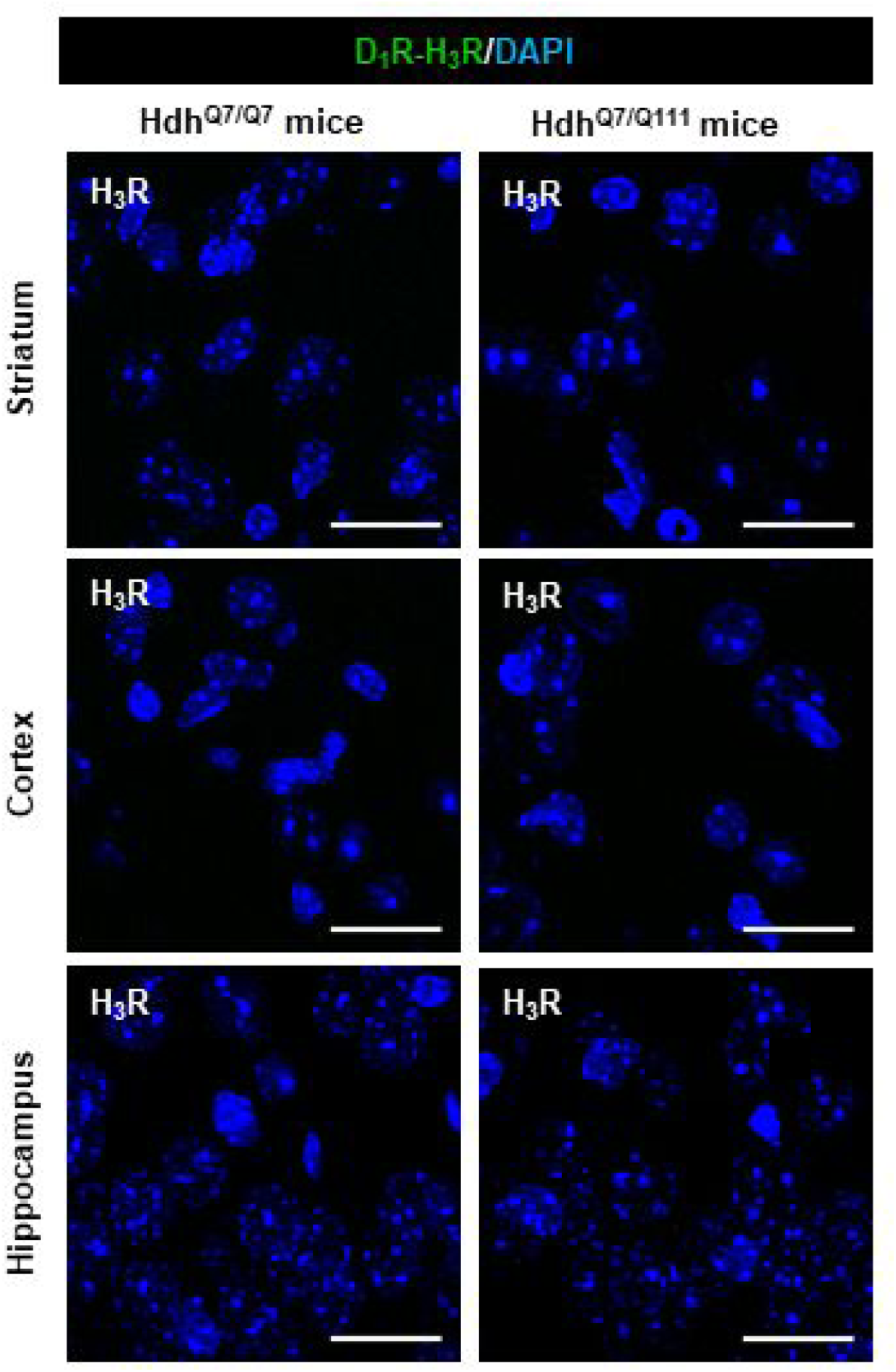
Negative controls for Proximity Ligation Assays (PLA) in mouse brain slices. Proximity Ligation Assays (PLA) were performed in the absence of the primary antibody against D_1_R, using striatal, cortical or hippocampal slices from 4-month-old Hdh^Q7/Q7^ and Hdh^Q7/Q111^ mice. In all slices, a lack of green spots around blue colored DAPI stained nucleus was observed. Scale bar: 20 μm.

**Figure S8.**
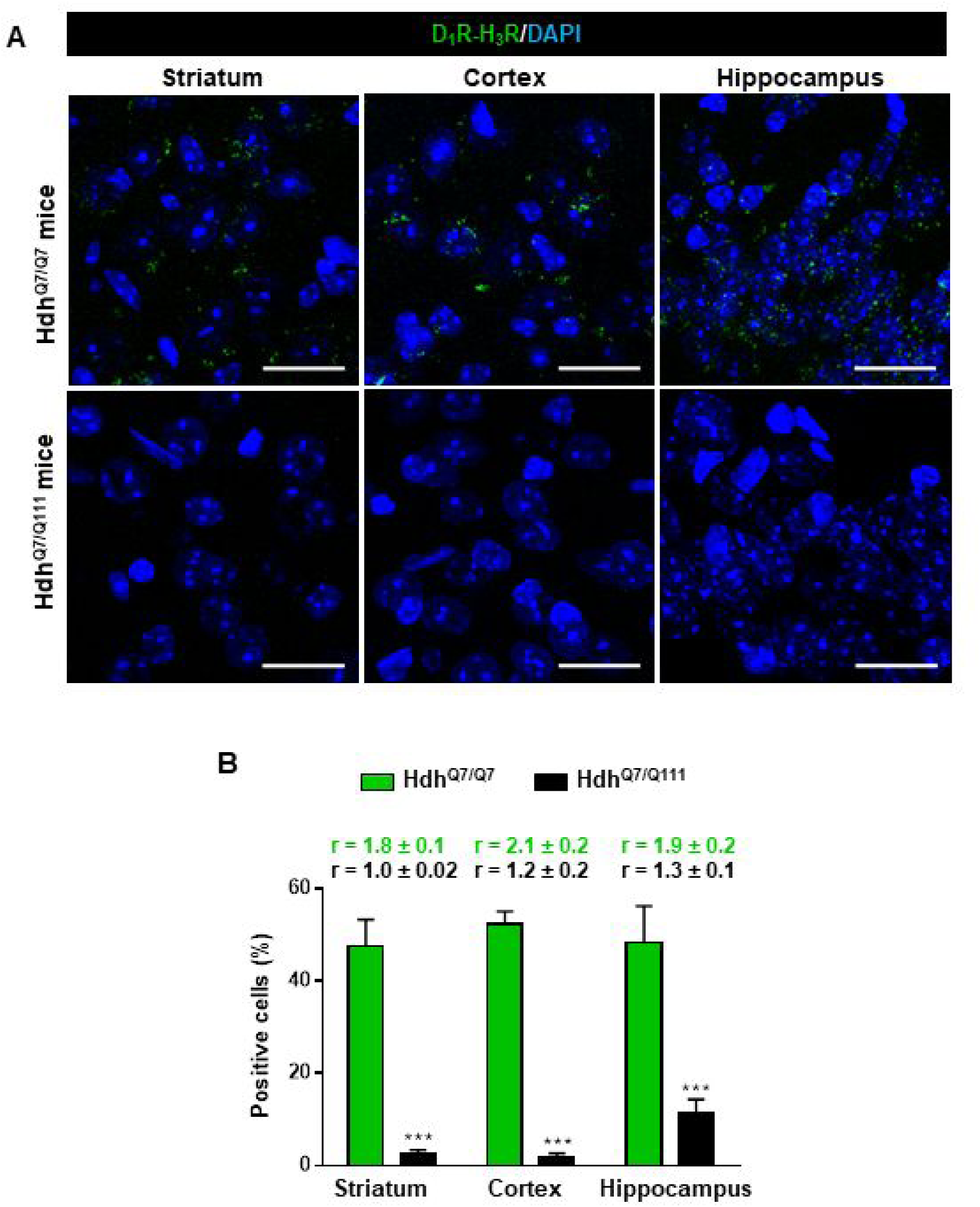
Expression of D_1_R-H_3_R heteromers in 6-month-old Hdh^Q7/Q7^ and Hdh^Q7/Q111^ mice chronically treated with saline. In (**A**), Proximity Ligation Assays (PLA) were performed in striatal, cortical and hippocampal slices from 6-month-old Hdh^Q7/Q7^ and Hdh^Q7/Q111^ mice treated with saline. D_1_R-H_3_R heteromers were visualized as green spots around blue colored DAPI stained nucleus in Hdh^Q7/Q7^ mice but not in Hdh^Q7/Q111^ mice chronically treated with saline. Scale bar: 20 μm. In (**B**), the number of cells containing one or more green spots is expressed as the percentage of the total number of cells (blue nucleus). *r values* (number of green spots/cell containing spots) are shown above each bar. Data (% of positive cells or r) are the mean ± SEM of counts in 600-800 cells from 4-8 different fields from 3 different animals. Student’s *t* test showed significant differences in D_1_R-H_3_R heteromer expression (***p<0.001) compared to the respective Hdh^Q7/Q7^ mice.

**Figure S9.**
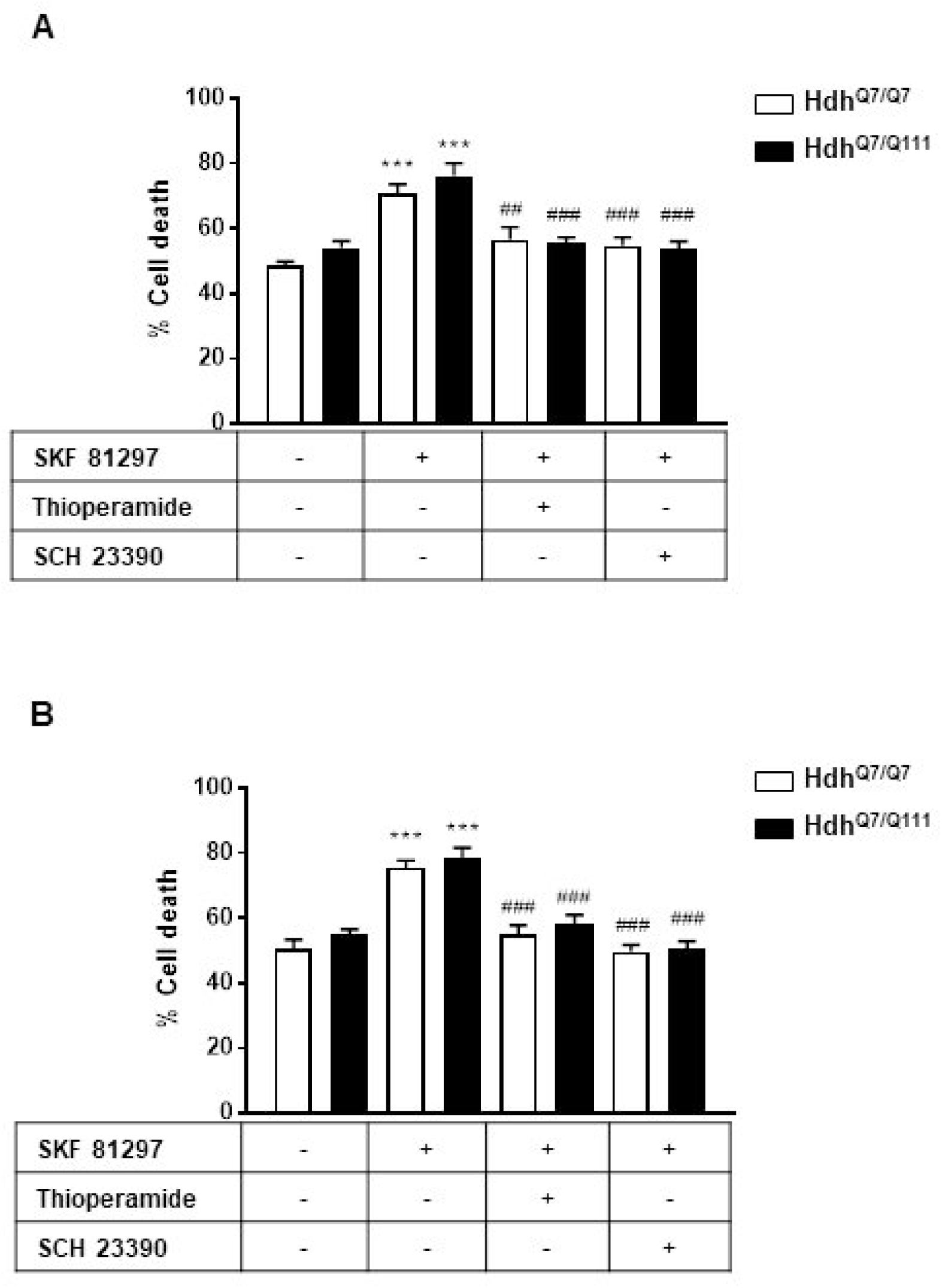
Functional D1R-H3R heteromers are expressed in 5-month-old HdhQ7/Q7 and HdhQ7/Q111 mice. Striatal (A, C) and cortical (B, D) organotypic slice cultures from 5-month-old HdhQ7/Q7 and HdhQ7/Q111 mice were pre-treated for 60 min with vehicle, H3R antagonist thioperamide (10 μM) or VUF5681 (10 μM) (C and D) or D1R antagonist SCH 23390 (10 μM) (A and B) before the addition of SKF 81297 (50 μM) and after 48h cell death was determined. Values represent mean ± SEM (n = 5 to 8) of percentage of cell death. One-way ANOVA followed by Bonferroni post hoc tests showed a significant effect over vehicle treatment (***p < 0.001) or over SKF 81297 treated slices (###p < 0.001).

**Figure S10.**
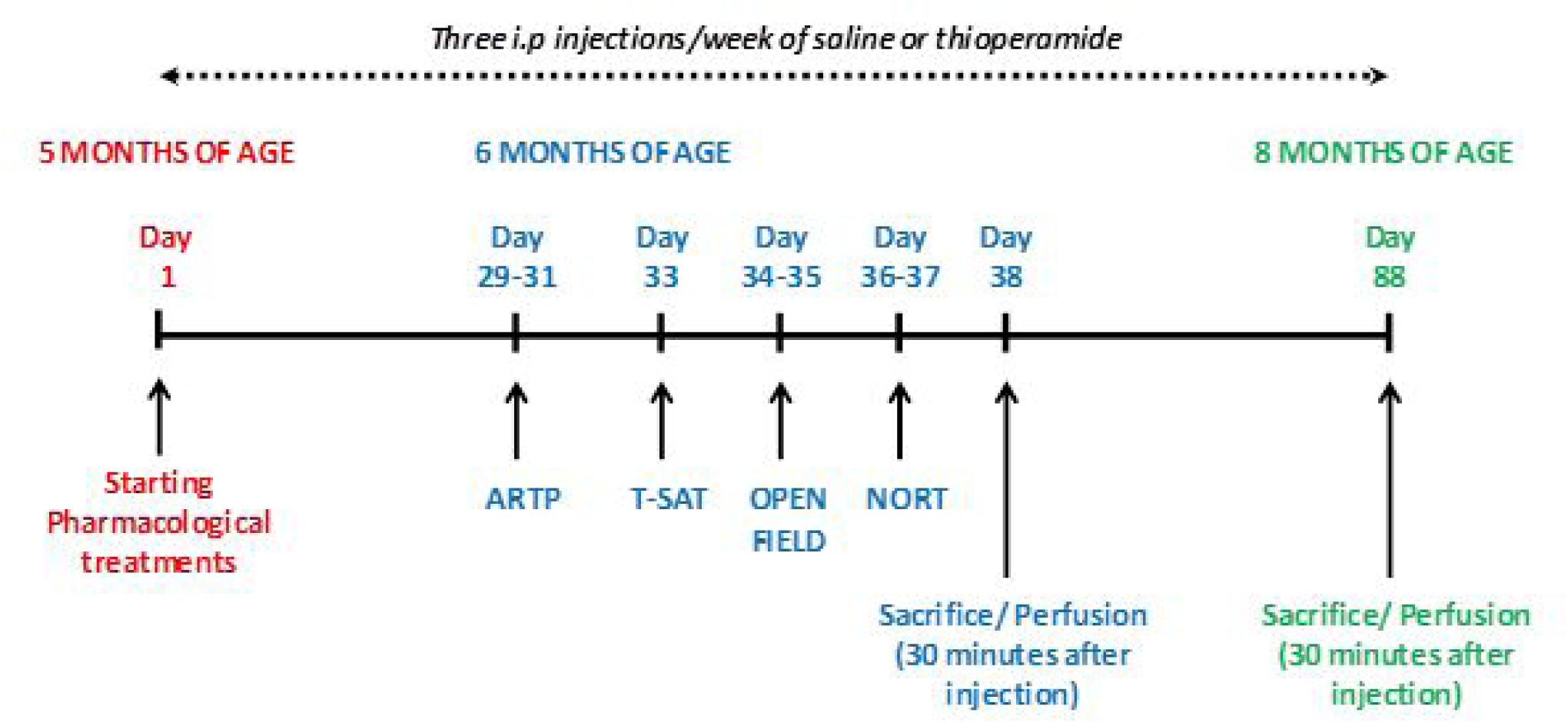
Schematic representation of pharmacological treatments and behavioral analysis performed after chronic treatment with saline or thioperamide. Three intraperitoneal injections per week of saline (NaCl 0.9% saline) or thioperamide (10 mg/Kg) were performed from 5-month-old to 8-month-old mice when the animals were sacrificed and perfused. Behavioral assessment started at 6 months of age with the evaluation of the ARTP, T-SAT, Open field and NORT. One cohort of animals was sacrificed and perfused 30 min after the last injection to evaluate PLA at 6 months of age. A second cohort of animals was sacrifice and perfused 30 minutes after the last injection to evaluate PLA at 8 months of age.

**Figure S11.**
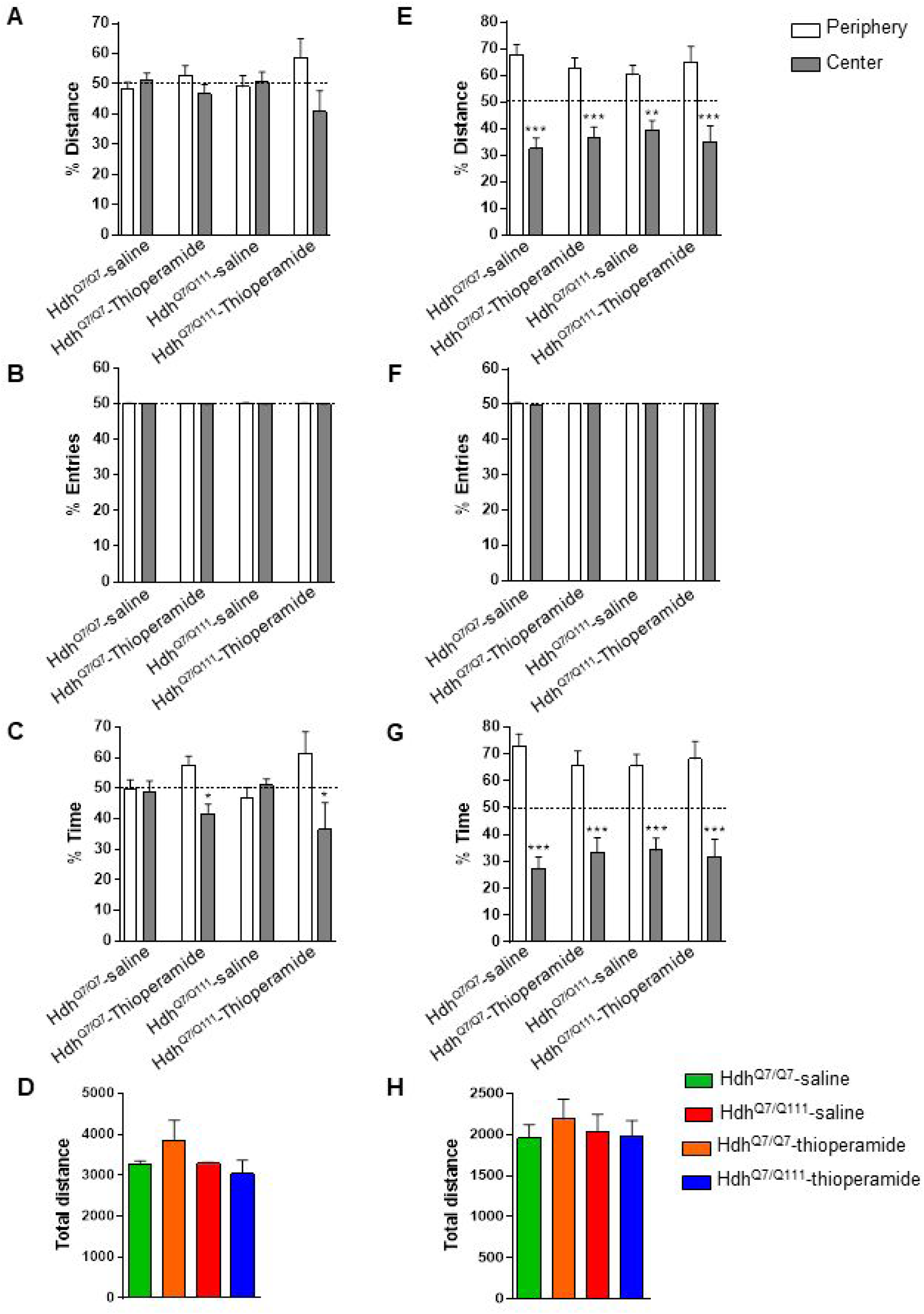
No significant differences in the open field habituation were found between treatments and genotypes. Motivation and anxiety differences between genotypes and treatments were analyzed by measuring the percentage of distance (**A** and **E**), the percentage of entries (**B** and **F**) and the percentage of time (**C** and **G**) between the periphery and the center in the open field arena at the first (**A**, **B** and **C**) or second (**E**, **F** and **G**) day of habituation in the open field arena. The spontaneous locomotor activity differences between genotypes and treatments were analyzed by measuring the total distance rove for each animal at first (**D**) or second (**H**) day of habituation in the open field arena. After two days of habituation in the open field arena, all mice behave equal. Data represents mean ± SEM. Statistical analysis was performed using one-way ANOVA with Bonferroni *post hoc* comparisons; *p<0.05, ***p<0.001 compared to the periphery.

**Figure S12.**
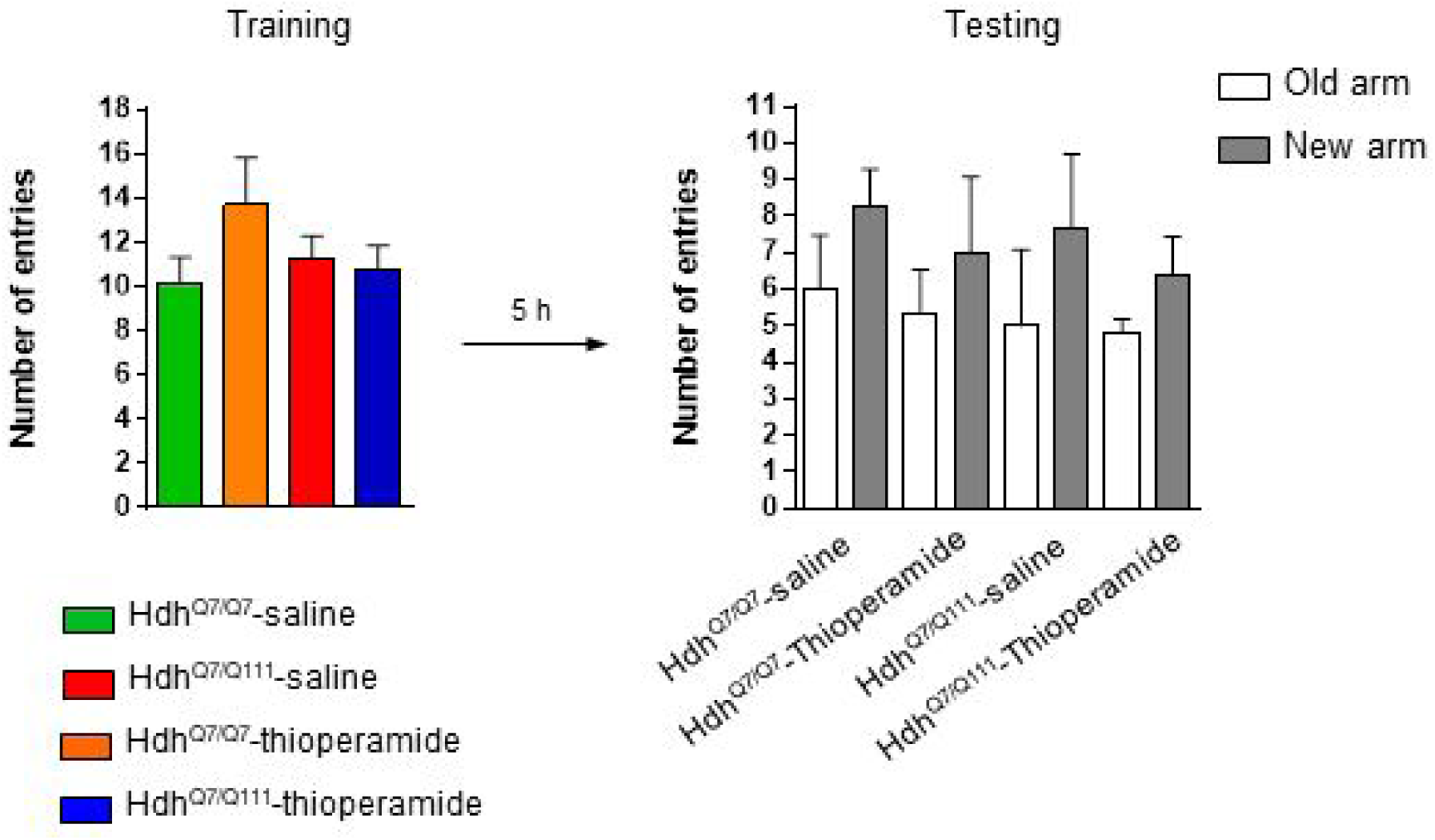
Training session in the T-SAT showed similar number of arm entries in all genotypes and treatments. 6-month-old Hdh^Q7/Q7^ and Hdh^Q7/Q111^ mice following the injection protocol in S8 showed no differences in spontaneous locomotor activity or anxiogenic components in training sessions of the T-maze. 11 saline-treated Hdh^Q7/Q7^ mice, 10 thioperamide-treated Hdh^Q7/Q7^ mice, 7 saline-treated Hdh^Q7/Q111^ mice and 9 thioperamide-treated Hdh^Q7/Q111^ mice were evaluated. Data represents mean ± SEM.

**Figure S13.**
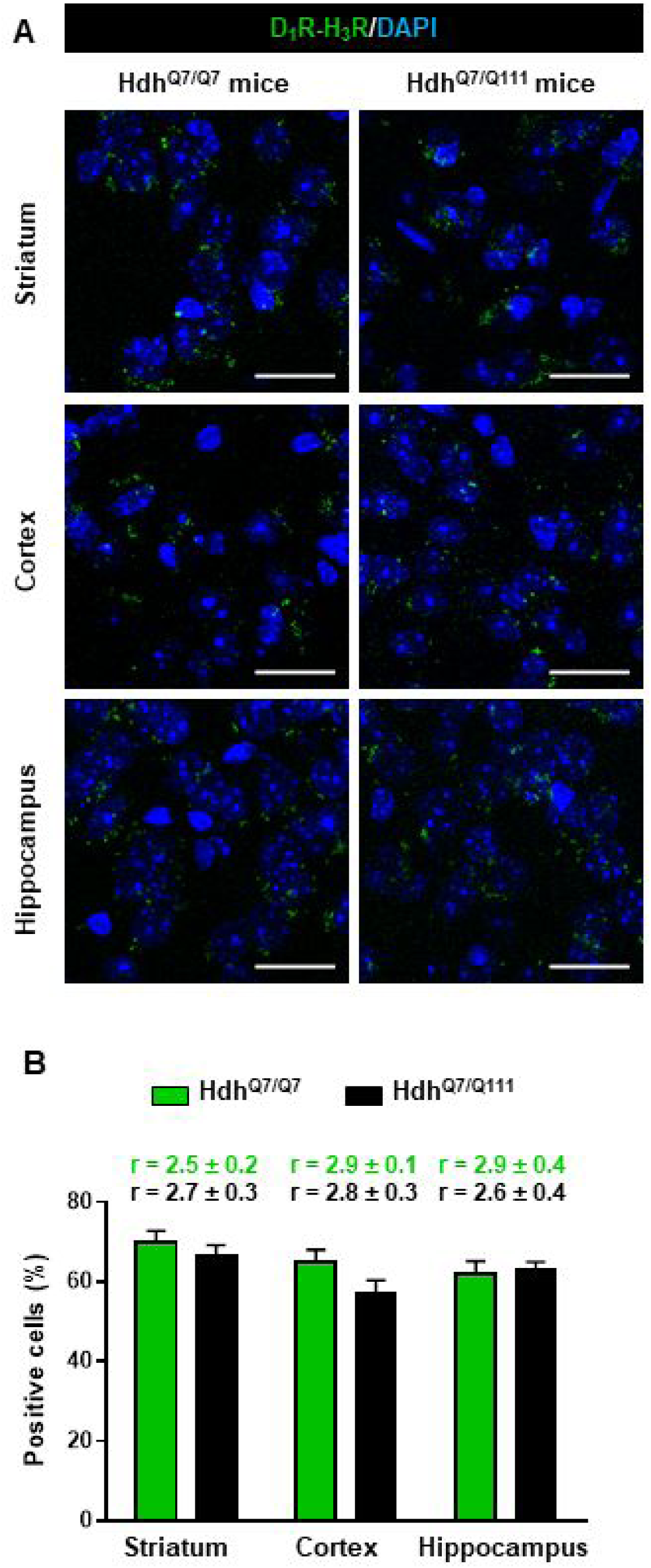
Expression of D_1_R-H_3_R heteromers in 8-month-old Hdh^Q7/Q7^ and Hdh^Q7/Q111^ mice chronically treated with thioperamide. In (**A**), Proximity Ligation Assays (PLA), were performed in striatal, cortical and hippocampal slices. D_1_R-H_3_R heteromers were visualized as green spots around blue colored DAPI stained nucleus in 8-month-old Hdh^Q7/Q7^ and Hdh^Q7/Q111^ mice chronically treated with thioperamide. Scale bar: 20 μm. In (**B**), the number of cells containing one or more green spots is expressed as the percentage of the total number of cells (blue nucleus). *r values* (number of green spots/cell containing spots) are shown above each bar. Data (% of positive cells or r) are the mean ± SEM of counts in 600-800 cells from 4-8 different fields from 3 different animals. Student’s *t* test showed no significant differences in D_1_R-H_3_R heteromer expression in thioperamide-treated Hdh^Q7/Q111^ mice compared to the respective Hdh^Q7/Q7^ mice.

**Figure S14.**
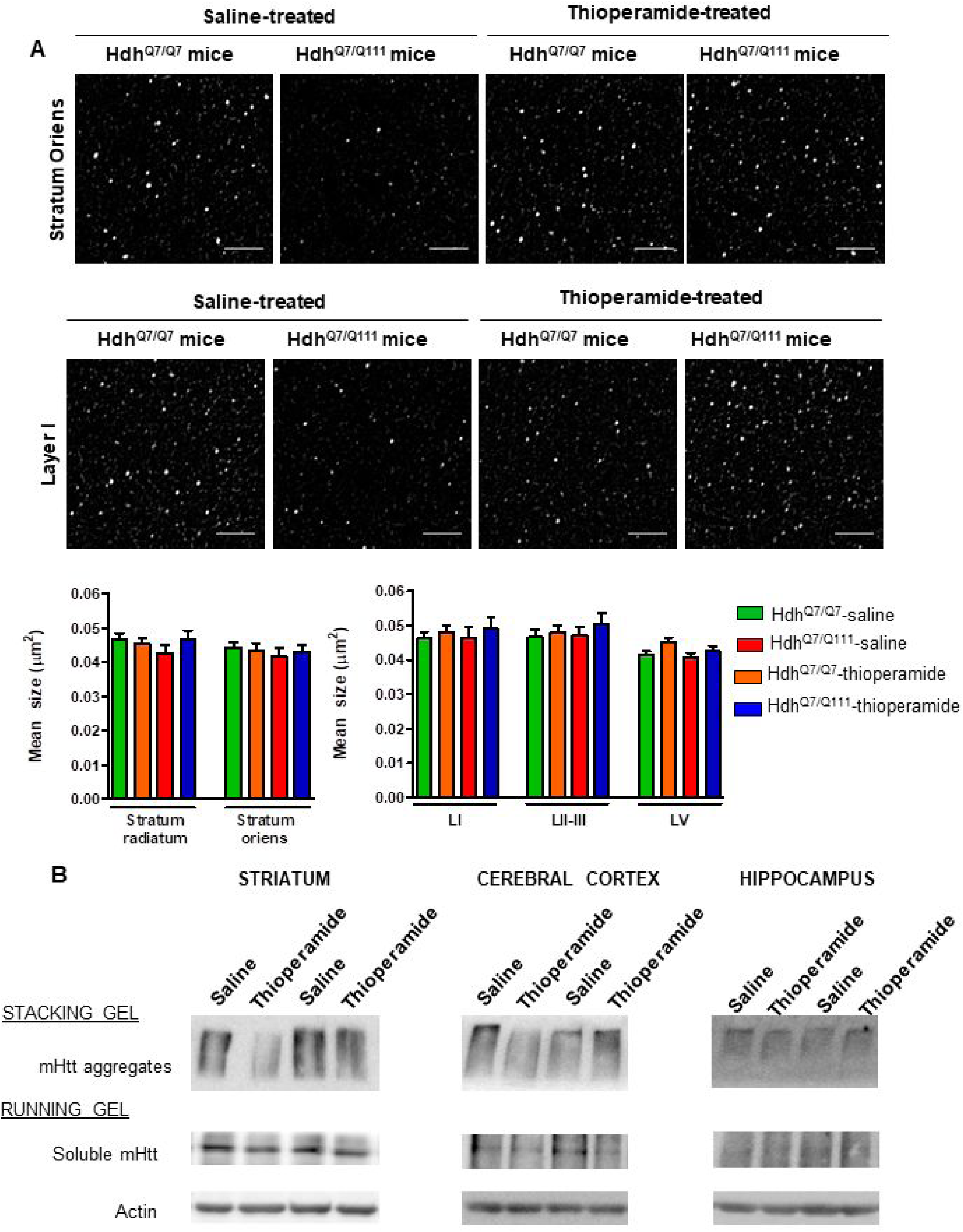
Biochemical and Pathological Effects of Thioperamide treatment. In (**A**), representative images showing spinophilin-immunoreactive puncta in the *stratum oriens* of CA1 hippocampus and in layer I of motor cortex area 1 (M1) of saline and thioperamide-treated WT Hdh^Q7/Q7^ and knock-in Hdh^Q7/Q111^ mice at 6 months of age. Quantitative analysis of the mean size of spinophilin-immunoreactive puncta in the *stratum radiatum* and *stratum oriens* of CA1 hippocampus and in layers I, II-III and V of motor cortex area 1 (M1) are shown as mean ± SEM (n= 9 images from three animals/group). Statistical analysis was performed using Student’s two-tailed t test. No significant differences were found. Scale bar: 5 μm. In (**B**), Representative Western blots of total striatal, hippocampal and cortical extracts from 6-month-old saline and thioperamide-treated knock-in Hdh^Q7/Q111^ mice. The blots were probed with 1C2 antibody for mutant huntingtin (mHtt). In samples from both saline and thioperamide-treated Hdh^Q7/Q111^ mice insoluble oligomeric forms of mHtt were detected in the stacking gel and soluble forms were detected in the running gel.

**Figure S15.**
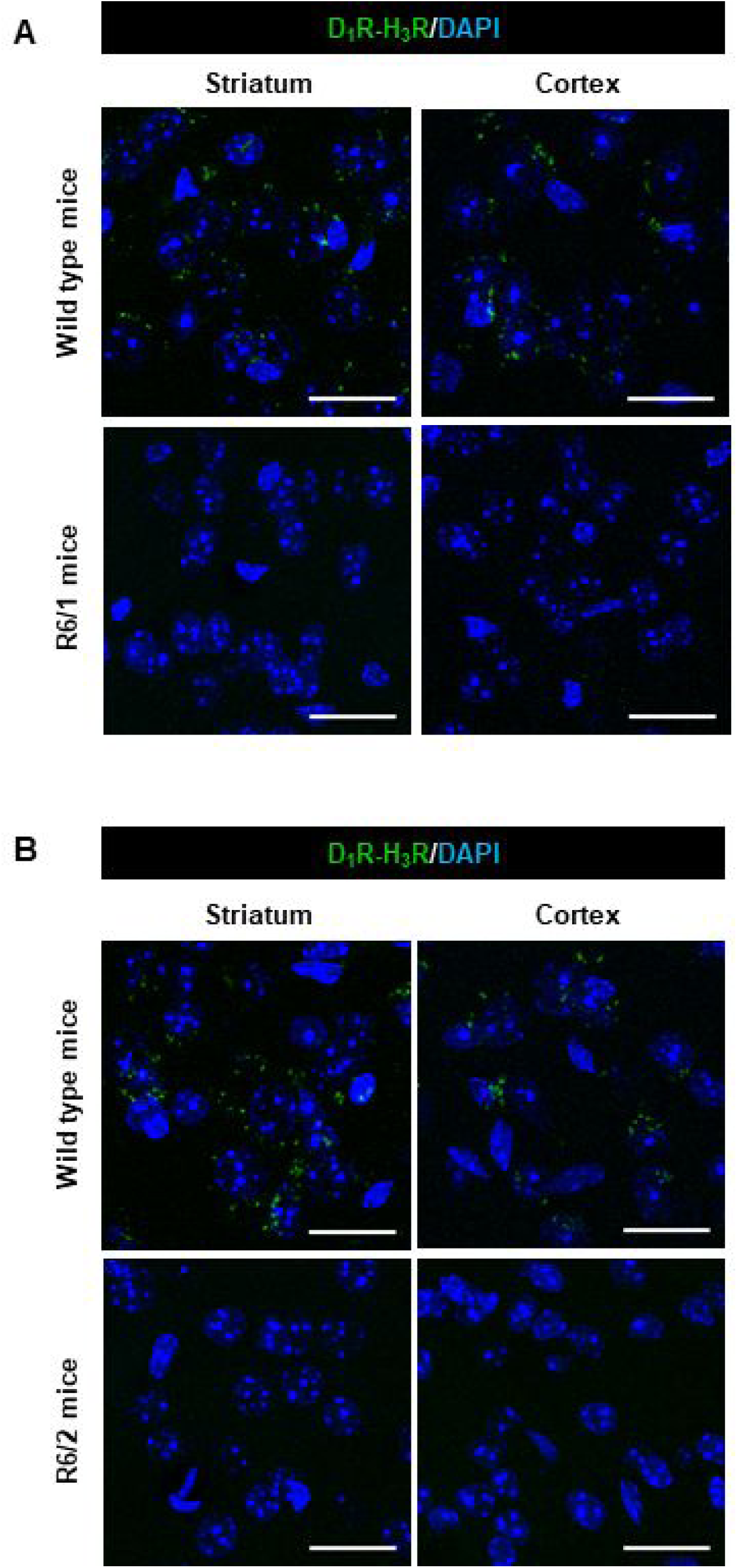
D_1_R-H_3_R heteromer are not expressed in HD R6/1 and R6/2 mouse models. Proximity Ligation Assays (PLA) were performed using striatal or cortical slices from age matched wild type littermates (WT) and 4-month-old R6/1 (**A**) or 8-week-old R6/2 mice (**B**). D_1_R-H_3_R heteromers were visualized only in wild-type mouse slices as green spots around blue colored DAPI stained nucleus. Scale bar: 20 μm.

